# Single-cell profiling of cortical tubers in tuberous sclerosis complex shows molecular structure preservation and massive reorganization of metabolism

**DOI:** 10.1101/2024.10.31.621014

**Authors:** Frederik Nørby Friis Sørensen, Tin Luka Petanjek, Mirte Scheper, Rasmus Rydbirk, Irina Korshunova, Jasper Anink, Angelika Mühlebner, James D. Mills, Zdravko Petanjek, Eleonora Aronica, Konstantin Khodosevich

## Abstract

Tuberous sclerosis complex (TSC) is a multisystemic genetic disorder associated with loss-of-function mutations in the *TSC1* or *TSC2* gene, which lead mTOR pathway hyperactivation and epileptogenesis. Cortical tubers are the hallmark of TSC and represent disorganized cortical structure underlying the generation of focal seizures. Here, we report single-nucleus RNA sequencing in resected cortical tubers vs matched pediatric controls. Strikingly, in spite of severe cortical disorganization, we found that cortical tubers preserve all neuronal subtypes, even the rarest ones. Moreover, we showed that principal neurons largely preserve spatial position based on transcriptional signatures. Principal neurons and layer 1-2 GABAergic neurons that modulate upper cortical circuits exhibited the largest gene expression changes. Interestingly, multiple mTOR pathway gene expression changes in TSC counteracted mTOR hyperactivation. TSC neuronal, but not glial, networks exhibited massive metabolic reorganization with a reduction in mitochondrial respiration and a concomitant switch to fatty acid metabolism. Finally, we show that neuron-specific AMPA receptor signaling might underlie epileptogenesis in TSC and could represent a potential candidate for therapeutic targeting.

## Introduction

Tuberous sclerosis complex, or simply tuberous sclerosis (TSC), is a rare autosomal dominant genetic disorder that affects several major organs of the body, including the brain^1–3^. It is characterized by monogenic loss-of-function mutations in either the *TSC1* or *TSC2* gene^4^. TSC can become manifest in various conditions in the brain and often most severely in treatment-resistant epilepsy and autism spectrum disorder (ASD)^5,6^. The TSC1/2 complex is part of the canonical mTORC1 pathway and inhibits mTOR (mechanistic Target of Rapamycin) signaling by inhibiting the mTOR activator RHEB (Ras homolog enriched in brain)^7^. Activation of the mTOR pathway leads to protein synthesis and increases metabolism, enhancing synthesis of nucleotides and lipids and activating glycolysis; inversely, catabolic pathways such as autophagy and lysosome biogenesis are inhibited^8^. Inhibition by TSC1/2 counteracts mTOR activation and restricts cell growth. In the brain, the balance between mTOR and TSC1/2 coordinates multiple developmental processes, such as neuronal maturation and axonal growth^9^. In TSC, the epileptic phenotype might arise already within the first year after birth^10^, when developmental processes are peaking, which is likely due to lack of TSC-mediated mTOR inhibition.

The etiology of epileptogenesis in TSC is still poorly understood. Hallmark pathological characteristics of TSC include cortical tubers with cytomegalic neurons, reactive astrocytes, and altered synaptogenesis^4,6,11,12^. Typical cortical layer structure within cortical tubers is lost and disorganized^1,4^. The lack of cortical structure and miswiring of neuronal networks within the epileptic focus are thought to be the main contributors to epileptogenesis, with perturbed developmental processes playing the key role in epileptogenic network miswiring^2^.

Studies on both animal models and resected tubers from patients have provided important knowledge about the morphology, circuits, and molecular hallmarks of TSC^2^ and identified potential developmental mechanisms, such as impaired neuronal migration and maturation^13,14^. Yet, both human iPSC-derived neurons and animal models cannot recapitulate the pathology of the disease entirely, especially regarding the high complexity of cortical neuronal types that are present in the human brain^15,16^.

An alternative approach to determine how neuronal complexity is perturbed in the epileptic focus and might contribute to epileptogenesis would be to study resections of brain tissue from epilepsy surgery using single-cell/nucleus RNA sequencing (sc/snRNA-seq). The high potential for this technology to reveal disease-related mechanisms in complex human brain disorders has been shown by multiple studies investigating various brain disorders, such as schizophrenia^17,18^, autism^19^, Alzheimer’s disease^20^, and multiple sclerosis^21^. For epilepsy, such an approach was first applied to analyze cortical epileptogenic regions associated with temporal lobe epilepsy (TLE)^22^. The data unraveled previously unknown proepileptogenic mechanisms, such as increased action of AMPA auxiliary subunits and large metabolic and developmental changes in molecular signaling, and identified deep-layer principal neurons, PVALB and VIP GABAergic neurons, as the most vulnerable neuronal subtypes potentially forming a network underlying epileptogenesis^22^. Other sc/snRNA-seq analyses revealed a glial role in such epileptogenic networks^23^.

Therefore, we initiated an in-depth, unbiased study to characterize the transcriptional landscape, cellular identity, and potential epileptogenic networks of frontal cortical tubers of TSC using snRNA-seq on resected TSC surgical samples matched to control pediatric samples. Using comprehensive computational analyses, we discovered that, despite a highly disorganized cortical layer structure, virtually all layer-specific neuronal types are present in cortical tubers, even very rare ones. Furthermore, using single-molecule FISH (smFISH), we demonstrated that molecular signatures associated with layer specificity were preserved although, histologically, a cortical lamination structure was lacking. Interestingly, the largest gene expression changes appeared in principal neurons of L2_3_CUX2 and L5_6_FEZF2 together with ID2 GABAergic neurons, the latter residing mainly in L1 and targeting upper cortical circuits and apical dendrites of L5_6_FEZF2 neurons^24,25^. Although cell type-specific molecular patterns are maintained in TSC, the metabolism-associated transcriptomic landscape was extensively modified, exhibiting a strong reduction in the generation of mitochondrial energy and a concomitant increase in fatty acid metabolism. Interestingly, in spite of mTOR hyperactivation at protein level, at transcriptional level we identified potential homeostatic mechanisms counteracting mTOR hyperactivation and circuit overexcitation. Finally, we found neuron-specific changes in AMPA receptor signaling, including AMPA auxiliary subunits, which might provide targets for future therapeutics.

## Results

### All neuronal subtypes of the neurotypical cortex are present in cortical tubers of TSC

Cortical tubers resected from the frontal cortex of pediatric patients carrying *TSC2* mutations and corresponding postmortem neurotypical control samples (11 and 6 individuals, respectively, ranging between 2 months and 17 years of age, Extended Data Table 1) were processed for snRNA-seq (Fig. 1a). To ensure that the postmortem gene expression signature had only limited impact, only very low postmortem interval (PMI) samples were used (6-9 h, Extended Data Table 1). Mutations of the *TSC2* gene in TSC samples were previously identified^26,27^, and a total of nine point mutations were represented among TSC2 samples (Fig. 1b): Six were frameshift mutations (deletions or insertions) and three were missense/nonsense mutations causing either faulty protein readout or termination of translation. Regardless of the mutation location site, all were loss of function.

**Figure 1.**
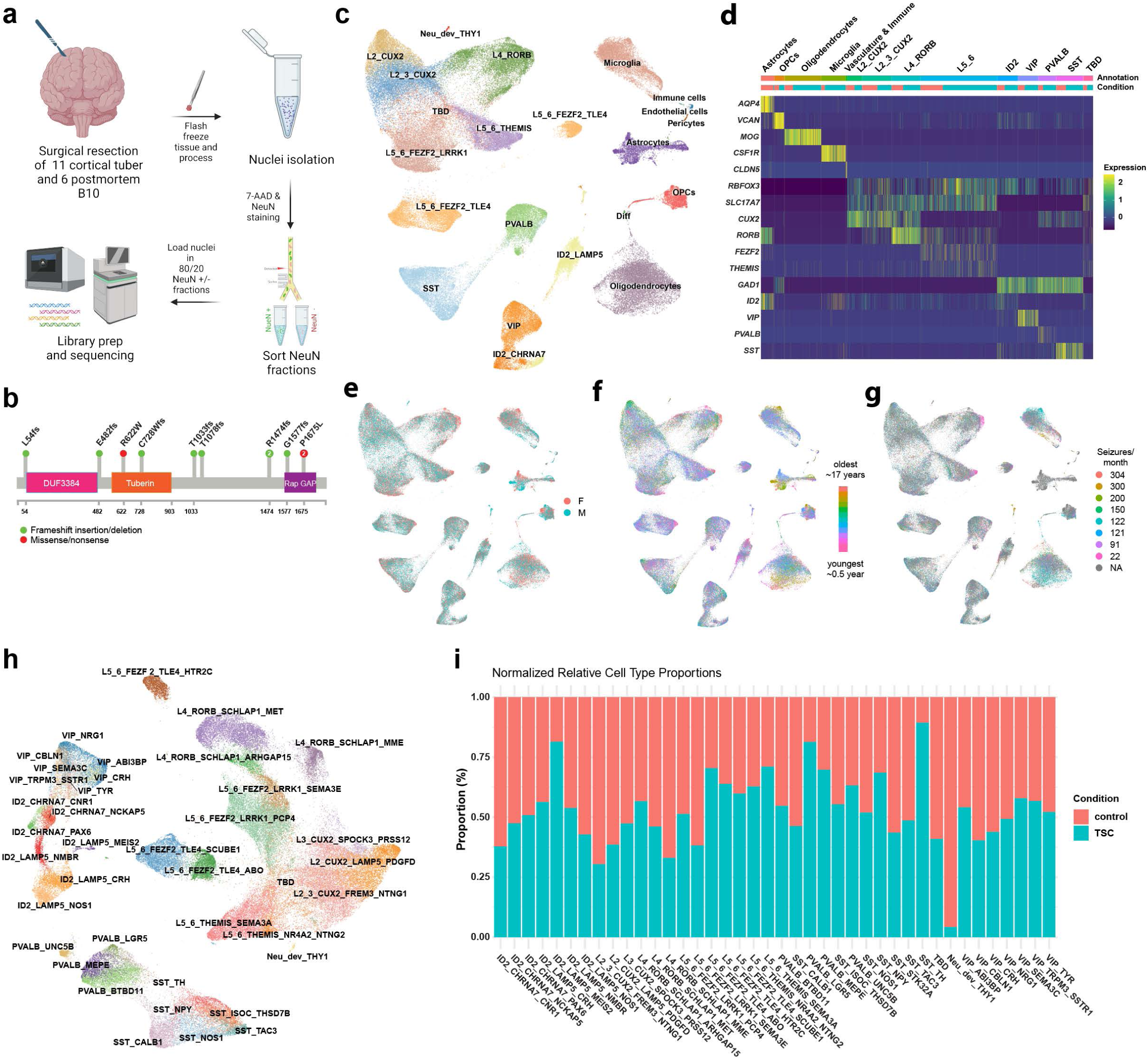
Single-nucleus RNA-sequencing of TSC and neurotypical control samples. (a) Experimental scheme. (b) List and type of TSC2 mutations in the samples used in the study. (c) UMAP representation of the analyzed nuclei, colored by cell type. (d) Expression of cell-type markers (note that Diff and Neu_dev_THY1 have too few cells to be shown on the heatmap). (e) - (g) UMAP representation of the major covariates in the dataset, colored by sex (e), age (f), and seizure frequency (g). (h) UMAP representation of principal excitatory and GABAergic inhibitory neuronal subtypes. (i) Distribution of each neuronal subtype in control and TSC samples.

The sample cohort was thoroughly balanced for potential confounders, such as sex and age (Extended Data Table 1). During tissue processing each sample was sorted using general nucleus marker 7-AAD followed by neuronal nucleus marker NeuN. To ensure that snRNA-seq captured a sufficient number of neurons and to avoid overrepresentation of oligodendrocytes (see Methods), NeuN+ and NeuN-fractions were collected and mixed at a ratio of 80% to 20% to perform snRNA-seq using the 10x Chromium v3.1 assay. In total, 166,524 nuclei were identified after sequencing using Cellranger (45,944 mean reads per nucleus) and after secondary cell calling with Cellbender and quality control filtering, which included ambient RNA, low-quality cells, cells with high mitochondrial reads, and doublets (Extended Data Fig. 1a-f, 2a-d, 3a-f), 100,282 nuclei (64,322 from TSC and 35,960 from controls) passed for further analysis. The resulting nuclei were of high quality with 8,445 median unique molecular identifiers and 3,017 median genes detected. The efficiency of this sorting strategy was confirmed after the sequencing with approximately 80% neurons and 20% non-neuronal cells identified in the snRNA-seq data (Extended Data Table 1).

Cell types were annotated manually using a combination of literature-based markers and previously analyzed datasets^17,22,28^ in a hierarchical manner, initially dividing them into glial cells (astrocytes, oligodendrocytes, oligodendrocyte progenitor cells [OPCs], and microglia), neurons (excitatory neurons, GABAergic inhibitory neurons), immune cells, and cells of vascular origin (pericytes and endothelial cells) (Fig. 1c,d, for non-neuronal cells Extended Data Fig. 4a-h, for neuronal cells Extended Data Fig. 5a-h). Neurons were subdivided further using layer-specific markers for the excitatory neurons L2-3_CUX2, L4_RORB, L5-6_FEZF2, and L5-6_THEMIS and markers of the cardinal families of GABAergic inhibitory neurons PVALB, SST, VIP, and ID2 (Fig. 1c,d). Importantly, biological covariates, such as sex, age at resection, or seizure frequency, had limited impact on snRNA-seq data clustering (Fig. 1e-g). Final subclustering of neurons into individual subtypes revealed 25 inhibitory GABAergic and 15 excitatory neuronal subtypes (Fig. 1h,i, Extended Data Fig. 5a-h). One excitatory cell population expressing a mixture of canonical layer markers was named “to be determined” (TBD). Although we confirmed that these cells were not doublets and they passed-low quality filtering (Extended Data Fig. 5e,f), we could not annotate this population. Owing to this and as the TBD cluster consists of cells with a generally lower transcriptional depth than the rest (Extended Data Fig. 5g,h), we excluded the TBD cluster from subsequent differential analyses.

Surprisingly, although the classical laminar-wise neuronal positioning in cortical tubers was found to be severely disorganized in neuroanatomical studies^1,4,29,30^, at the molecular level all canonical cell types and all neuronal subtypes that are present in neurotypical cortex were also present in TSC tubers (Fig. 1h,i, Extended Data Fig. 4a). Even the rarest excitatory and GABAergic neuronal subtypes, such as L5_6_THEMIS_NR4A2_NTNG2 (0.84% of total neurons) and L3_CUX2_SPOCK3_PRSS12 (1.10% of total), and ID2_LAMP5_MEIS2 (0.23% of total) and SST_TH (0.27% of total), were present in TSC tissue (Fig. 1i). Thus, our data demonstrate the advantage of molecular phenotyping at the single-cell level for identifying specific neuronal subtypes in the disorganized cortex.

### Deep-layer excitatory neurons, PVALB inhibitory neurons, and microglia are enriched in TSC

Although, as shown above, TSC contains all neuronal and glial subtypes that are identified in the normal cortex, the cellular abundance of specific cell subtypes and their gene expression signatures in TSC might be altered, which might underlie epileptogenesis. Thus, we performed differential analysis of cellular composition and gene expression between conditions. Rigorous statistical testing showed no significant impact of covariates on gene expression or cell composition differences, including sex, age, snRNA-seq processing steps and sample loading on flowcells (Extended Data Fig. 6a-h). By projecting overall gene expression or cellular abundance on multidimensional scaling (MDS) plots^17,31^, we showed that two conditions were clearly separated, indicating that TSC has unique transcriptomic and cell composition signatures (Fig. 2a).

**Figure 2.**
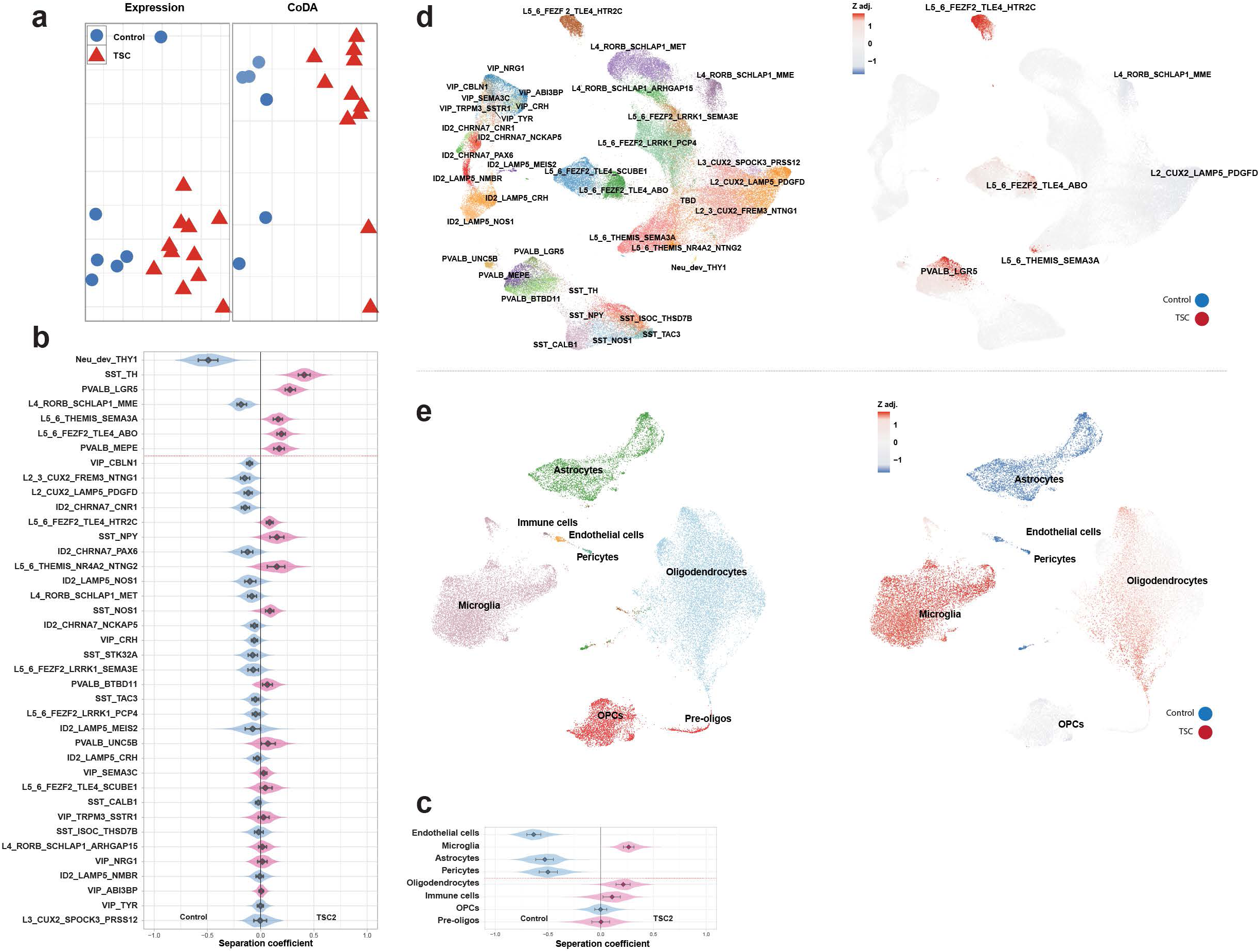
Subgrouping of TSC and control samples and cellular composition changes in TSC. (a) Multidimensional scaling plot visualizing similarities for gene expression (left) and cellular composition between all samples analyzed by snRNA-seq. Note clear division in control and TSC subgroups. One control sample (top position in the left plot) is derived from the youngest subject and thus shows some differences. (b) Compositional differences for neuronal subtypes between TSC and control groups analyzed by Compositional Data Analysis (CoDA). Separation coefficient (x-axis) represents the scale of changes, with the left half for increase in controls and right half for increase in TSC. Cell subtypes above the red line are significantly different, as adjusted p values <0.05 (analyzed by the Benjamini-Hochberg method). (c) Same as (b) for non-neuronal subtypes. (d) Changes in cellular composition between TSC and control groups analyzed using UMAP embedding. Left: UMAP embedding colored by subtype of neurons, right: Z adjusted score for cell density differences (see Methods). (e) Same as (d) for non-neuronal subtypes.

To characterize changes in cell-type composition in TSC, we implemented Compositional Data Analysis (CoDA) as implemented in *Cacoa*^17,31^. While simply comparing proportions of cell types makes them inherently interdependent on each other, CoDA corrects for such interdependency, thus allowing for independent comparison of cellular abundance for each cell type in a sc/snRNA-seq dataset. We analyzed the cellular composition of NeuN+ and NeuN-cells separately to avoid introducing bias due to the enforced 80% to 20% mixing ratio of NeuN+ to NeuN-cells. Among excitatory neurons, layer-4 subtype L4_RORB_SCHLAP1_MME was decreased in TSC, whereas deep-layer excitatory neurons L5_6 THEMIS_SEMA3A and FEZF2_TLE2_ABO were enriched in TSC (Fig. 2b). Additionally, the Neu_dev_THY1 neuronal subtype was enriched in controls -- this subtype was detected almost exclusively in the youngest neonatal control sample (age 2 months) and expressed excitatory, but not GABAergic inhibitory markers (Extended Data Fig. 7). Interestingly, several subtypes of GABAergic inhibitory neurons were enriched in TSC, including major subtypes of the PVALB neurons PVALB_MEPE and PVALB_LGR5, where the former corresponds to interlaminar/translaminar basket cells and the latter likely representing another basket-cell subtype^32^. Additionally, SST_TH neurons were the subtype of GABAergic neurons with the highest enrichment in TSC (Fig. 2b). SST_TH neurons are a rare subtype of GABAergic neurons that have been reported to express TH protein and are thus potentially dopaminergic neurons^33^.

To circumvent annotation bias, we performed cluster-free compositional analysis, i.e., by identifying changes in cell composition using graph-based density on UMAP^17,31^. This analysis confirmed an increased cellular abundance of deep-layer excitatory neurons (L5_6_FEZF2_TLE4_HTR2C, L5_6_FEZF2_TLE4_ABO and L5_6_THEMIS_SEMA3A) and inhibitory PVALB_LGR5 neurons in TSC (Fig. 2d), as above. Additionally, lower densities of upper middle-layer excitatory neurons (L2_CUX2_LAMP5_PDGFD and L4_RORB_SCHLAP1_MME) were observed (Fig. 2d), which might indicate that TSC has the opposite impact on upper- and deeper-layer neurons by expanding deeper layers in TSC with concomitant reduction in the upper layer neurons.

An isolated look at non-neuronal cells shows strong compositional changes in several cell types (Fig. 2c). Microglia exhibited a large enrichment in TSC (Fig. 2c), thus confirming previous studies showing potential infiltration of microglia into cortical tubers and increased microglia activation in drug-resistant seizure patients^34–36^. However, we did not find evidence for inflammatory pathway upregulation in microglia in TSC (Extended Data Table 2). Other non-neuronal cell types were reduced, including both major blood vessel cell types – endothelial cells and pericytes (Fig. 2c), suggesting a perturbation of the blood-brain-barrier in TSC. Additionally, astrocytes were also reduced in TSC samples (both assessed by independent comparison of cellular abundances by CoDA – Fig. 2c, and graph-based density – Fig. 2e), which was unexpected, as astroglia have been described in TSC tissue both by morphological studies and immunophenotyping^37,38^. Finally, cluster-free analysis of non-neuronal cells confirmed all of our findings and, in addition, allowed us to detect enrichment of oligodendrocytes in TSC (Fig. 2e). This enrichment was likely missed in annotated cluster-based comparison (Fig. 2c) due to only part of the oligodendrocyte cluster being enriched in TSC (Fig. 2e). The finding suggests the existence of a specific oligodendrocyte population that is enriched in TSC.

### Massive gene expression changes in TSC that are associated with energy metabolism and glutamate signaling

To identify the molecular mechanisms underlying TSC etiology that can contribute to seizure generation within the epileptic focus, we assessed changes in gene expression for each cell type in TSC. First, to evaluate overall gene expression change per cell type, we implemented an expression distance metric^17,31^. Strikingly, we found that almost all subtypes of cells showed significant changes in gene expression in TSC compared to neurotypical controls (Fig. 3a,b), which suggests that there is a general, extensive change in gene expression across many cell types in TSC. Among neurons, the largest changes occurred in subtypes of excitatory neurons of L2_3 and L5_6 as well as in GABAergic inhibitory neurons of the ID2 family that mainly reside in L1 (Fig. 3a). The largest expression shifts in non-neuronal cell types were observed in astrocytes and oligodendrocyte precursor cells (OPCs) (Fig. 3b). The population sizes of other cell types such as endothelial cells, pericytes, and immune cells were too small to calculate the expression distance metric.

**Figure 3.**
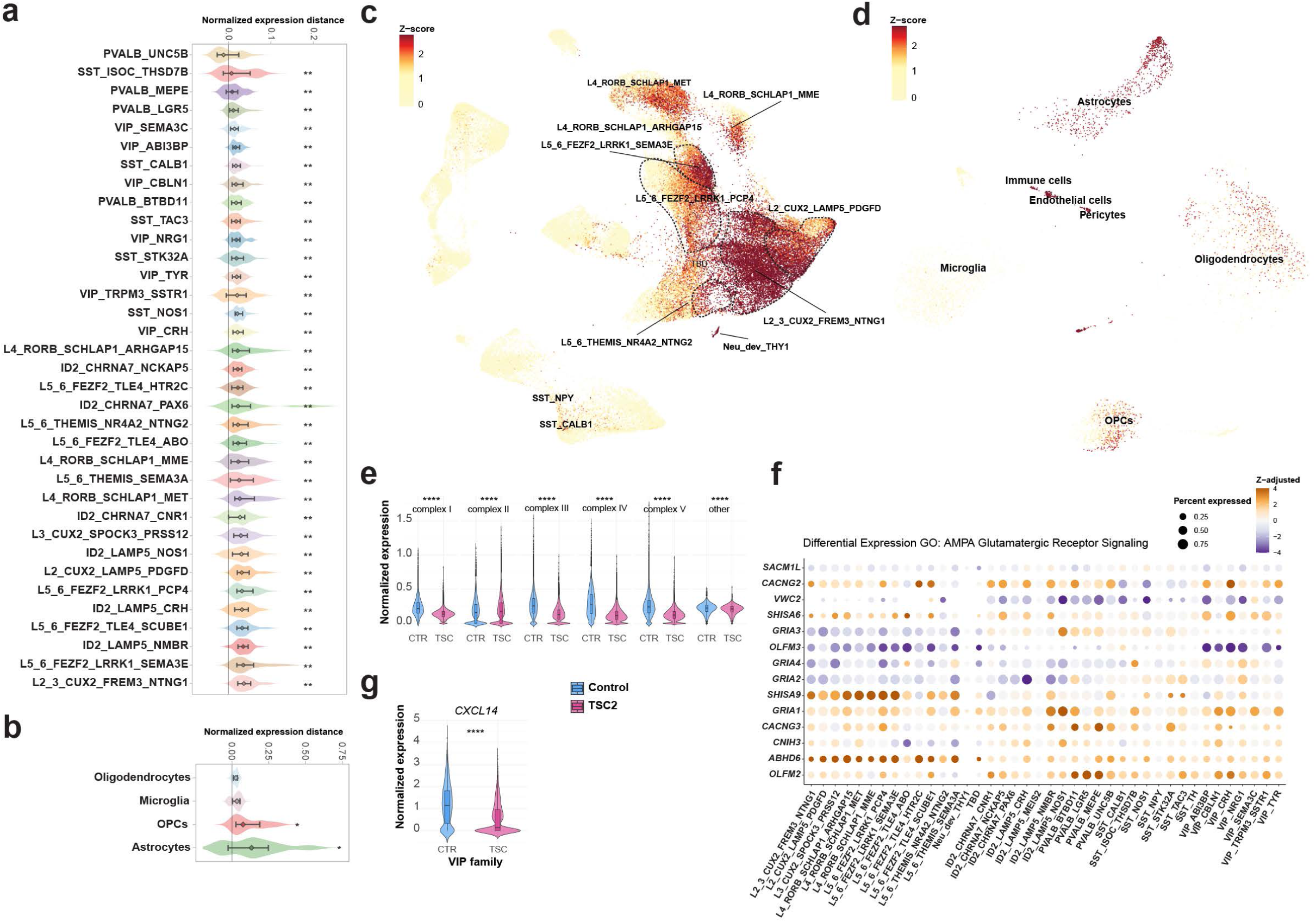
Gene expression changes in neuronal and non-neuronal subtypes in TSC. (a) Violin plots showing the change in transcriptome between TSC and control samples for each neuronal subtype as normalized expression distance metric. The distance is calculated based on Pearson linear correlation coefficient, normalized by the medium variation within the control and TSC2 groups. The subtypes are ordered based on the magnitude of expression distance change from lowest (top) to highest (bottom). Multiple correction was done by Benjamini-Hochberg. \*\**P* = 0.001 to 0.01. (b) Same as (a) for non-neuronal subtypes (metric for some subtypes with too few cells could not be measured). (c) Cluster-free analysis of gene expression changes in TSC neurons. The UMAP shows the extent of gene expression changes measured by adjusted p-value statistical significance levels and converted to Z score. (d) Same as (c) for non-neuronal subtypes. (e) Violin plots demonstrating changes in expression for genes coding for proteins involved in mitochondrial respiration. Separate violin plots for each complex are shown. “Other” category represents average expression of all genes that are involved in mitochondrial respiration, but not a part of complexes I-V. Bars show median with standard deviations; dots label the outliers. (f) Dot-plot representing expression levels of DE genes from GO term “AMPA receptor signaling” in TSC neuronal subtypes. Subtypes are ordered by principal neurons followed by GABAergic neurons. The size of the dots reflects the percent of cells within a subtype that expresses a gene. Orange and blue colors indicate up- and downregulation, respectively, and the intensity of the color corresponds to Z-adjusted value for expression change. (g) Violin plot showing changes in expression levels of *CXCL14* gene averaged for VIP neuronal family. Bars show median with standard deviations; dots label the outliers.

Cluster-free differential expression analysis can identify transcriptomic changes in cell populations that do not depend on UMAP clustering and are not linked to a specific cluster^17,31^. Interestingly, implementation of cluster-free differential expression analysis showed major changes in excitatory neuron subtypes across all layers, with the largest changes in upper L2_3_CUX2 neurons (Fig. 3c). This cluster-free approach also uncovered profound gene expression changes in blood vessel cells – endothelial cells and pericytes (Fig. 3d) – that could not be analyzed by classical cluster-based differential expression as there were too few of these cell types in the dataset. In addition, the cluster-free approach confirmed transcriptomic changes in OPCs and astrocytes. Taking both cluster-based and cluster-free comparisons of gene expression together, the most pronounced gene expression changes were in excitatory neurons across all layers (with L2_3 dominating), layer 1 GABAergic neurons of ID2 family and OPCs, astrocytes, and blood vessel cells.

To detect differentially expressed (DE) genes that can be associated with molecular changes in TSC, we performed DE analysis for each subtype. Interestingly, the genes involved in mitochondrial respiration (e.g., *UQCRB*, *ATP5F1E*, and *COX7C*) and protein biogenesis (e.g., ribosomal protein genes, *EEF1A1*, and *NACA*) were greatly downregulated across TSC neuronal subtypes (Extended Data Fig. 8,9, Extended Data Table 3). By assigning genes to complex I-V, we showed that downregulation affected all five complexes of the electron transport chain (Fig. 3e, Extended Data Fig. 10). Interestingly, we found a significant modulation of glutamate receptor signaling (Extended Data Fig. 11) and, in particular, of AMPA receptor signaling (Fig. 3f) in TSC neurons across multiple cortical layers, which resembles the data for TLE^22^. Across glutamate receptor genes, prominent changes were observed in kainate receptor genes: upregulation of *GRIK4* (principal neurons across all layers and several layer 1 ID2 subtypes of GABAergic neurons) and *GRIK5* (principal neurons across all layers), which was accompanied by downregulation of kainate auxiliary subunit gene *NETO2* specifically in L2_3_CUX2 principal neurons (Extended Data Fig. 11). While AMPA receptor subunits exhibited minor downregulation (*GRIA2*, *GRIA3*, and *GRIA4*) or upregulation (*GRIA1*) in principal neurons, several AMPA auxiliary subunits were profoundly upregulated, mainly in principal neuron subtypes, such as CKAMP44 (gene *SHISA9*), CKAMP52 (gene *SHISA6*), GSG1L (*GSG1L*), TARPg2 (*CACNG2*), and TARPg3 (*CACNG3*) (Fig. 3f).

Particularly strong upregulation was observed for the gene *SHISA9,* coding for the protein CKAMP44, mainly in L2_3_CUX2, L4_RORB, and L5_6_FEZF2 families of principal neurons (Fig. 3f). Among other DE genes, upregulation of the chemokine gene *CXCL14* in VIP neurons might indicate their morphogenesis (Fig. 3g, Extended Data Figure 12).

### Neuronal cells in TSC show profound downregulation of mitochondrial respiration pathways with a concomitant switch to fatty acids in energy generation

To infer the molecular mechanisms underlying epileptogenesis, we identified signaling pathways from DE genes using gene-set enrichment analysis (GSEA). We detected a profound downregulation of pathways related to protein/ribosomal biogenesis and energy generation across most neuronal and glial cell types (Fig. 4a,b). Interestingly, while a major energy generation process such as mitochondrial oxidative phosphorylation was downregulated (Fig. 4a,b, Extended Data Table 4), fatty acid-associated energy generation pathways were upregulated in TSC, in particular in excitatory neurons (Fig. 4a,b). A focused analysis on energy generation pathways further confirmed a massive decrease in oxidative phosphorylation gene expression with fatty acid usage being concomitantly upregulated (Fig. 4c), whereas there was only minor downregulation of glycolysis (Extended Data Table 5). This finding suggests that there is a potential switch in energy generation pathways to fatty acids, with the most pronounced switch in excitatory neurons of L2_3, L4, and L5_6 identities, as well as in GABAergic neurons of the VIP and PVALB families (Fig. 4c). The downregulation observed in mitochondrial respiration and protein/ribosomal biogenesis (Fig. 4a) are the opposite of what would be expected from the overactivation of mTOR^8,39^ and might indicate homeostatic processes at transcriptomic level counteracting mTOR hyperactivation at protein level. Nevertheless, these changes might indicate that maturation of neuronal circuits is impaired in TSC tubers since increased expression of protein/ribosomal biogenesis is required for axonal growth, and these pathways decline with maturation^40^.

**Figure 4.**
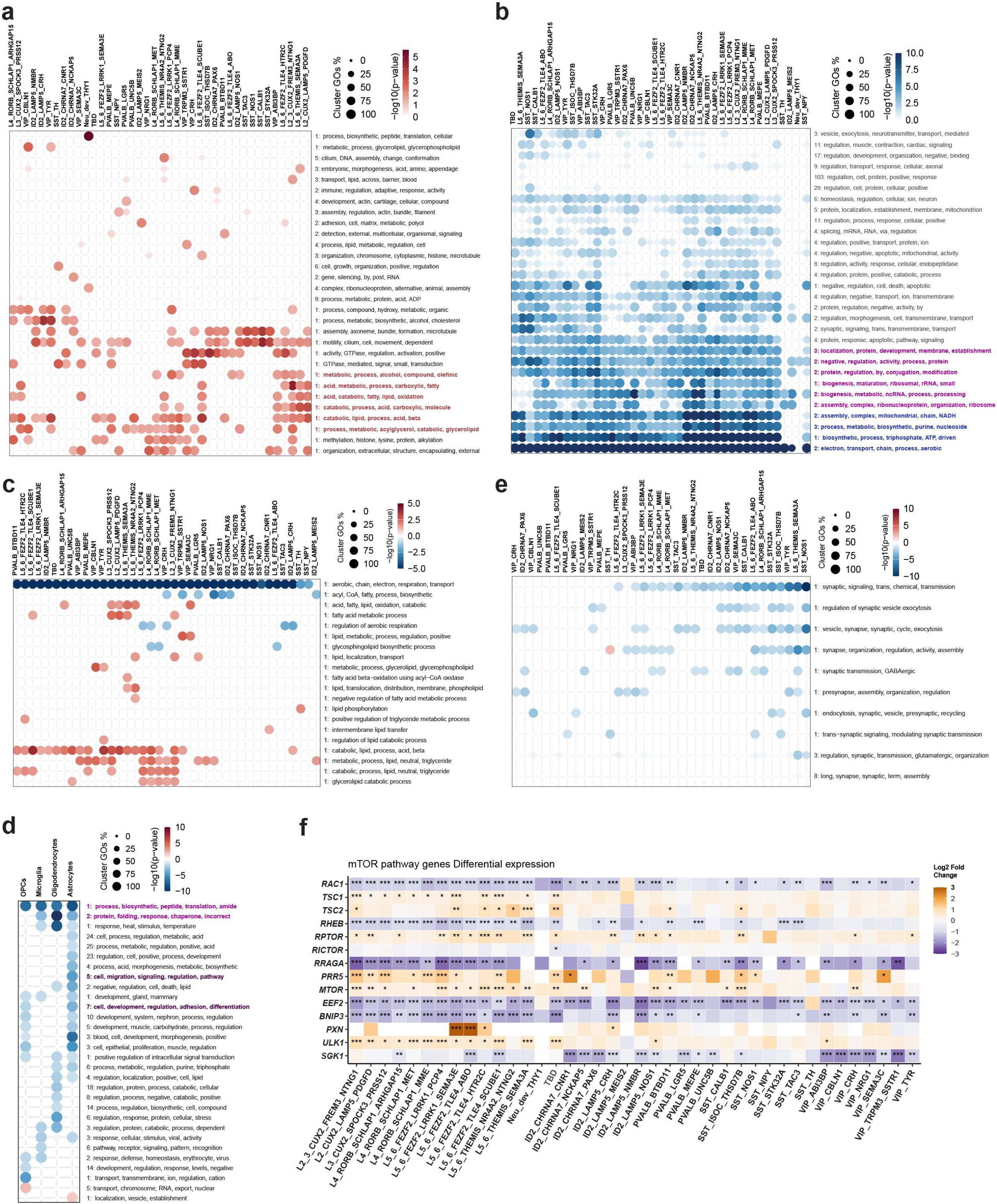
Differential pathway analysis for neuronal and non-neuronal subtypes in TSC. (a) GSEA for upregulated signaling pathways in neuronal subtypes in TSC. GO terms on the right are collapsed based on similarity of words within the terms and the number in front of each collapsed GO term line indicates the number of specific GO terms that are collapsed. Significance levels were calculated as in *clusterProfiler* with a wrapper around *fgsea* with default parameters. Collapsed GO term lines labeled with orange color correspond to fatty acid metabolism. (b) Similar as (a) for downregulated signaling pathways. Collapsed GO term lines labeled with blue and magenta colors correspond to oxidative phosphorylation and protein biogenesis pathways, respectively. (c) Focused GSEA on major energy generation pathways - oxidative phosphorylation, glycolysis, and fatty acid metabolism. Analysis is performed similar to that described in (a). (d) Focused GSEA on synaptic pathways. Analysis is performed similar to that described in (a). (e) GSEA for up- and downregulated signaling pathways in glial subtypes in TSC. Collapsed GO term lines labeled with dark purple and magenta colors correspond to cell migration/adhesion and protein biogenesis pathways, respectively. Analysis is performed similar to that described in (a). (f) DE for selected mTOR pathway genes (see full list in Extended Data Fig. 14. Heatmap showing DE of genes from mTOR pathway in TSC neuronal subtypes. Genes are ranked from the top to the bottom based on how upstream/downstream they are in the mTOR signaling pathway. Orange/blue coloring represents the log2 fold magnitude of gene expression changes.

Interestingly, these metabolic changes were specific for neurons: none of the glial cell types showed either dampening of mitochondrial oxidative phosphorylation or a switch to fatty acid metabolism (Fig. 4d). Overall, most of modulated pathways in glial cells were downregulated (Fig. 4d). The largest changes in molecular pathways in glial cells pertained to protein translation, affecting all glial cell types. In addition, astrocytes exhibited more pathway changes than other glial cell types and included downregulation of gene ontology (GO) terms associated with cellular migration and adhesion (Fig. 4d), indicating reduced astrocyte motility.

We found two more types of molecular pathways that were consistently downregulated in TSC neurons – synaptic and developmental pathways (Fig. 4a,b). A closer look at the synaptic pathways showed the largest downregulation in subtypes of SST GABAergic neuron and principal neurons of L5_6 family (Fig. 4e). Within the developmental pathways, there were a number of genes coding for proteins that are involved in axonal development, including cytoskeleton assembly, axon guidance, and related transcription factor genes (Extended Data Fig. 13). Both synaptic and developmental pathway downregulation might indicate perturbed synaptic plasticity. Surprisingly, although there is a lack of clear cortical layering structure and severe neuronal displacement in TSC2 lesions, we found only subtle changes in neuronal migration signaling pathways (Extended Data Table 6).

Finally, as *TSC2* mutations should perturb the mTOR pathway, we determined the relationship between the aforementioned differential pathways and mTOR signaling. Interestingly, we found a rather contrasting pattern of gene expression changes for principal neurons vs GABAergic neurons. Thus, several key mTOR pathway genes were mildly upregulated in principal neurons, either broadly across the layers (e.g., *TSC1*, *MTOR*, and *ULK1*) or specifically in certain layers/sublayers (*AKT1* in L2_3_CUX2_FREM3_NTNG1, *PXN* in L5_6_FEZF2_LRRK1_SEMA3E and L5_6_FEZF2_TLE4_ABO, and *PLD1* in L4_RORB_SCHLAP1_ARHGAP15 and L4_RORB_SCHLAP1_MET), but not in GABAergic neurons (Fig. 4f, Extended Data Fig. 14). Concomitantly, some downstream effectors of mTOR signaling were downregulated across both principal neurons and GABAergic neurons, such as EEF2, CYCS, YWHAG, YWHAB, and YWHAH (Extended Data Fig. 14). Most importantly, a few key mTOR pathway activation genes were downregulated, mainly in principal neurons, e.g., RHEB and RRAGA (Fig. 4f). Thus, although the current model of epileptogenesis in TSC proposes hyperactivation of mTOR^41^, our data show the existence of homeostatic molecular processes that can potentially counteract mTOR hyperactivation at transcriptomic level, such as extensive dampening of the expression of protein biogenesis components across most neuronal subtypes (*RRAGA* as amino acid sensor and *EEF2* as translation factor), and strong downregulation of key mTOR complex component *RHEB* in principal neurons.

### Neurodevelopmental and epigenetic transcription factor networks are enriched in TSC neurons

To gain insight into the transcription factor (TF) landscape underlying TSC pathogenesis we used CollecTRI^42^, a curated database of transcriptional regulatory interactions which contain signed TF from multiple resources (Fig. 5a, Extended Table 7). The majority of TF networks were positively enriched across DE genes, with many networks deriving from neurodevelopmental TFs, including REST, ZNF331, GTF2IRD1, GLI3, ERF, CDX2, and SOX5 (Fig. 5a). The highest score was attributed to the REST network, which was enriched in all principal neurons and most GABAergic neurons (Fig. 5a). REST is a major regulator of neuronal differentiation^43^, which controls the epigenetic mechanisms regulating gene expression^44^. Importantly, other epigenetic regulation networks were enhanced in TSC neurons, such as DNMT3a and KDM5C (Fig. 5a), both of them having previously been associated with seizures^45,46^. Overall, since increased levels of REST were reported to contribute to epilepsy^47^, the REST TF network might be at the core of epileptogenesis by modulating epigenetic mechanisms that promote pro-epileptic gene expression in TSC neurons, potentially leading to partial aberrant neuronal differentiation.

**Figure 5.**
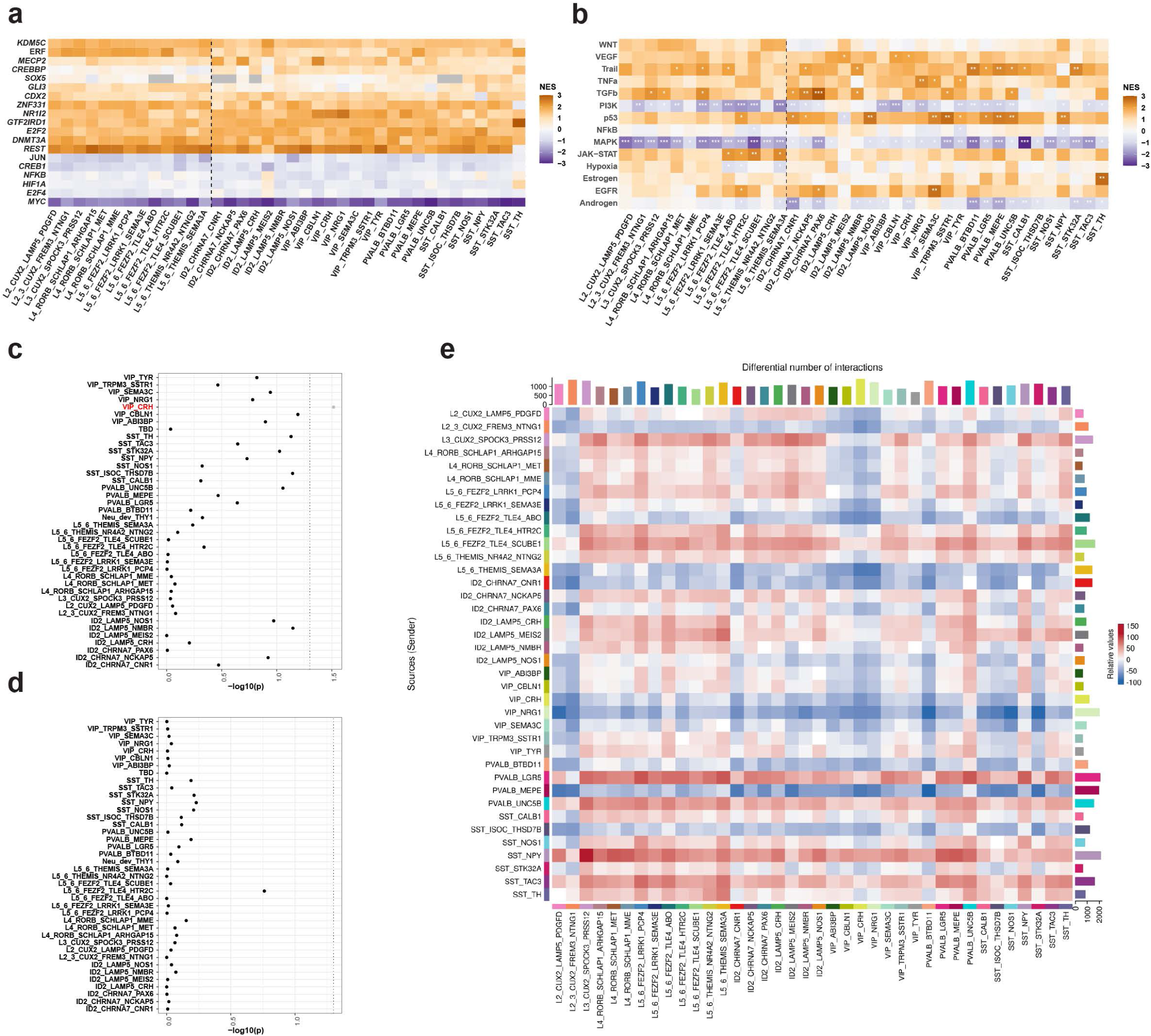
Enriched transcription factor networks and genetically associated DE genes in TSC. (a) Heatmap showing enriched transcription factor (TF) networks that were identified based the z score of DE genes, filtered for top 10 TF per cell type, scaled to -3,3 and presented by hierarchical clustering. Selected TF networks are shown; for full list see Extended Data Table 7. NES, Normalized enrichment score. (b) Hypergeometric test was used to identify overlap between DE genes in snRNA-seq TSC data with GWAS-linked genes in focal epilepsies (taken from^50^). Black dotted line indicates p-value cut-off. P-values were adjusted using Bonferroni correction. (c) Same as (b) for GWAS-linked genes in generalized epilepsies (taken from^50^). (e) Heatmap showing differential number of interactions between neuronal cell types. Sources of signaling are on the y-axis and receivers of the x-axis. The right and top colored bar represents the total sum of outgoing or incoming signaling in absolute values, respectively. Red and blue colors represent increased or decreased relative quantities of signaling between neuronal cell type pairs in TSC compared to control.

Among the negatively enriched TF networks, only one had a high score across most neuronal subtypes – the MYC network (Fig. 5a). Although MYC was not reported to be genetically associated with epilepsy, downregulation of MYC was associated with repetitive electroconvulsive seizures in a rat model^48^. Finally, many positively enriched TF networks showed the highest enrichment in specific types of neurons (Extended Data Fig. 15).

We used the above obtained regulons and inferred pathway activities based on prior knowledge using resources such as PROGENy (a model based on pathway signaling perturbation data)^49^; therefore, we performed an enrichment analysis for 14 curated pathways available at PROGENy. While many unrelated pathways displayed low significance, low enrichment or both, there was a strong negative enrichment for cell growth pathway activity, including MAPK (across principal neurons, PVALB, and SST GABAergic neurons) and PI3K (in L5_6 principal neurons) (Fig. 5b). These findings corroborate other analyses reported above showing that, despite that mTOR is disinhibited and subsequently hyperactivated due to loss-of-function TSC2, it does not lead to enhanced cell growth activation: on the contrary, cell growth-associated pathways are downregulated likely due to homeostatic mechanisms.

To gain further insight into epileptogenic mechanisms in TSC, we integrated our snRNA-seq DE genes with the largest genome-wide association study (GWAS) on epilepsy^50^. We collected GWAS summary data on generalized epilepsies and focal epilepsies and implemented the CELLECT/CELLEX workflow for summary statistics integration using MAGMA^51^. By integrating focal epilepsy GWAS into the data, we showed that GWAS-linked genes were enriched in TSC DE genes in GABAergic neuronal subtypes, although significance was reached only for the VIP_CRH subtype (Fig. 5c). Remarkably, this enrichment pattern was gone when integrating snRNA-seq with generalized epilepsy GWAS (Fig. 5d). Thus, integration with genetics confirmed that the DE genes we identified by snRNA-seq are specifically related to focal epilepsy (TSC).

Finally, to leverage potential changes in neuronal connectivity in TSC, we performed differential cell-cell interaction analysis in our snRNA-seq dataset using CellChat^52^. Both the number of interactions (Fig. 5e) and interaction strength (Extended Data Fig. 16) showed a similar differential pattern, which included several key features: (1) stronger interaction between principal neurons, with 2 blocks (L3_CUX2 to L5_6_FEZF2_LRRK1 and L5_6_FEZF2_TLE4 to L5_6_THEMIS_NR4A2), (2) weaker interaction of VIP GABAergic neurons with other neurons, (3) stronger interaction between ID2 GABAergic neurons and principal neurons, and (4) stronger interaction of PVALB_LGR5 and weaker interaction of PVALB_MEPE basket neurons. While (1) is likely related to increased excitation in epileptic circuits, (2)-(4) seem to be homeostatic mechanisms directed towards increasing inhibition in epileptic circuits, such as the increase in basket neuron inhibition on bodies of principal neurons, increase in ID2 neuron inhibition on dendrites of principal neurons, and decrease in disinhibition provided by VIP neurons.

### The layer-specific molecular signature of principal neuron subtypes is preserved in cortical tubers

We showed before that cortical tubers preserve all neuronal subtypes (Fig. 1i), and the changes in TSC arise due to perturbed gene expression that might underlie epileptogenesis, mainly metabolic and glutamate signaling pathways. However, another key question arises as to whether TSC subtypes also preserve at least some spatial position specificity, which is particularly important for translational perspectives and repair of epileptic circuits. Although a vast amount of previous data on the pathology of TSC shows disorganized cortical layers^1,29,30^, these data are based mainly on labeling neuronal bodies with NeuN antibody. Thus, we performed cross-labeling to determine the molecular signatures for multiple principal neuron subtypes, to detect their spatial position relative to each other, and to reveal whether they preserve at least partial organization into layers (Fig. 6a). We selected CUX2, RORB, and FEZF2 mRNAs in order to identify multiple subtypes of principal neurons based on single mRNA presence or overlap (Fig. 6b). We performed smFISH to identify these mRNA markers (Fig. 6c, d) and quantified the distribution of single- and double-labeled cells across the whole cortical column from the pia to white matter, separated in 7 equal bins (Fig. 6e-h).

**Figure 6.**
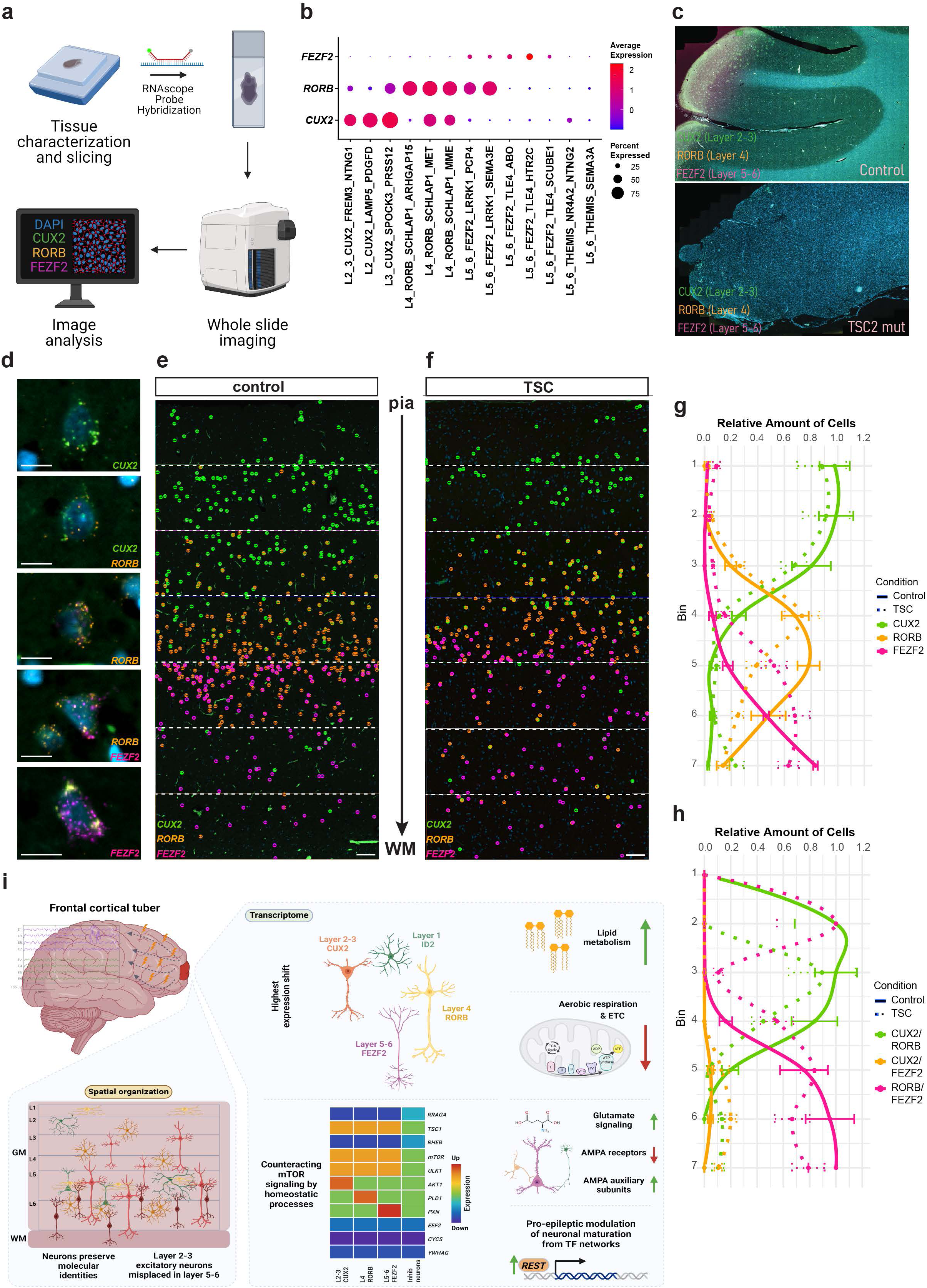
Spatial layer-wise molecular signature of principal neurons in TSC. (a) Experimental overview for smFISH labeling of cortical tissue from TSC patients and control subjects. (b) Dot plot showing distribution of *CUX2*, *RORB*, and *FEZF2* gene expression across all subtypes of principal neurons. (c) Overview of triple smFISH labeling showing gross level lack of gray vs white matter structure in TSC lesions. (d) Representative images showing single- and dual-labeled neurons. Scale bars are 10 μm. (e), (f) Pseudocoloring for molecular signatures of principal neuron subtypes: 1=CUX2, 3=RORB, and 4=FEZF2. Note that dual-marker neurons (CUX2/RORB, RORB/FEZF2, and CUX2/FEZF2) are labeled as overlay of both colors. For quantification, cortical columns were divided by 7 equal bins from the pia to white matter (WM). Scale bars are 100 μm. (g) Line plot showing relative amounts of *CUX2-*, *RORB-,* and *FEZF2*-positive cells across bins and condition. Points represent the observed relative amounts per bin with error bars showing standard deviation (SD). Smoothed lines calculated by LOESS method are used to show trends. Colors: *CUX2* (green), *RORB* (orange), and *FEZF2* (pink). Solid lines show control (n=5) and dotted TSC (n=5). (h) Line plot showing relative amounts of double-positive *CUX2:RORB*, *CUX2:FEZF2,* and *RORB*:*FEZF2* cells across bins and condition. Points represent the observed relative amounts per bin with error-bars showing standard deviation (SD). Smoothed lines calculated by LOESS method are used to show trends. Colors: *CUX2*:*RORB* (green), *CUX2*:*FEZF2* (orange), and *RORB*:*FEZF2* (pink). Solid lines show control (n=5) and dotted TSC (n=5).

Our most striking finding was that the overall spatial distribution of subtypes of principal neurons based on their molecular signatures was largely preserved in TSC (Fig. 6e-h). This was true for all the molecular signatures we used: CUX2, CUX2/RORB, RORB, RORB/FEZF2, CUX2/FEZF2, and FEZF2 (Fig. 6g,h). Nevertheless, there were several local changes, including an increase in lower layers for CUX2 principal neurons in TSC (bin 7, Fig. 6g) and a concomitant increase in FEZF2 principal neurons in the upper bins (bins 6-7 in controls vs 5-7 in TSC, Fig. 6g). For mixed marker subtypes, there was an increase in CUX2/FEZF2 colabeling in TSC, which was rather rare in controls (Fig. 6h), and RORB/FEZF2 subtype neurons were present in upper layers in TSC, which were absent in the corresponding layers of control cortex (bin 2, Fig. 6h). Thus, it appears that the cortical structure in TSC is largely preserved, while some specific populations are displaced from upper to lower layers or vice versa. The major changes pertain to metabolism reorganization and glutamate signaling, which might lead to changes in connectivity that contribute to the formation of epileptogenic networks in TSC (Fig. 6i).

## Discussion

TSC is a developmental epilepsy whose etiology is based on mutations in the *TSC1* or *TSC2* gene^4^. However, in spite of the clear association with mutated genes, it has been challenging to generate treatments specifically targeting TSC and ameliorating epileptogenesis. Thus, novel approaches are required to understand the molecular mechanisms underlying the generation of seizures in cortical tubers in TSC for diagnostics and drug target discovery. Here, we implemented snRNA-seq on a set of frontal TSC samples, all with *TSC2* loss-of-function mutations, and compared them with high-quality, low PMI control samples from the same cortical area. Our data revealed that, in spite of the fact that virtually no layer-wise structure could be identified by classical pathological assessments, cortical tubers contain all neuronal subtypes that preserve their subtype-specific molecular signatures, as in their counterparts in normal tissue. Moreover, we demonstrated that subtypes of principal neurons largely preserve their spatial layer-specific distribution based on molecular signatures and that the major shift was observed in L2_3_CUX2 neurons being misplaced towards L5-L6 and vice versa – L5_6_FEZF2 in L2-3. The largest gene expression changes exhibited principal neurons and L1_2_ID2 GABAergic neurons. Thus, together with misplacement of L2-3 neurons, it indicates potential impairment of upper cortical layer circuits that are evolutionary novel circuits in humans^24,53,54^. Mechanistically, we demonstrated a switch to fatty acid metabolism and proexcitation glutamate signaling in principal neurons, which could be important for developing therapeutic strategies.

As one of the key findings, we demonstrated that cortical tubers do preserve all subtypes of neurons, even very rare ones. Thus, although TSC1/2 proteins have a significant role in proliferation, differentiation, and cell death^4,8^, they do not induce radical changes in cell-type composition in cortical tubers. Furthermore, using smFISH, we revealed that cortical tubers preserve the layer-specific molecular signatures for principal neuron subtypes. This indicates that, although by classical histology cortical tubers do not show any layering, according to the transcriptional profiles, the layers are largely preserved, with a population of neurons being displaced.

Accordingly, important conclusions can be drawn for the field. First of all, our findings suggest that the etiology of TSC is based on specific postmitotic changes for several distinct types of neurons that might have increased vulnerability to TSC mutations, whereas large proliferation and differentiation changes reported as a potential mechanism in TSC^5,55^ might be partially balanced by homeostatic processes in human developing brain. The vulnerable neuronal populations include L2_3_CUX2, which are late-born principal neurons during the inside-out generation of human cortex^56^. Interestingly, our data suggest that late-born neurons and late cortical developmental processes might be the most affected and thus most relevant to TSC disease etiology. Hence, the increase in abundance of early-born L5_6_FEZF2/THEMIS neurons can be due to a delayed switch from early to late born cortical principal neurons. Likewise, increased abundance of L2_3_CUX2 neurons in lower layers and L5_6_FEZF2/THEMIS in upper layer can arise by impairment of neuroblast migration during late cortical development. This modulation of migration and differentiation is likely limited to the embryonic period, when both processes are particularly sensitive to neurodevelopmental risk factors^57–60^, including the impact of TSC1/2^61^. Nevertheless, active epileptogenesis might modulate terminal differentiation/maturation in neurons in childhood and adolescence since we found positively enriched TF networks controlling neuronal differentiation (e.g., REST^44,62^) in TSC.

Another prominent finding was that protein biogenesis and mitochondrial respiration genes were highly downregulated in TSC cells accompanied by a switch to fatty acid-dependent energy metabolism due to upregulation of corresponding gene expression pathways. Interestingly, the metabolic shift was specific to neurons and not glial cells. The switch to fatty acid-dependent energy metabolism together with downregulation of mitochondrial respiration might indicate that, due to epileptogenesis, neuronal networks may experience energy shortages and neurons try to adjust their energy demands by turning to fatty acids. Interestingly, this downregulation of protein biogenesis and mitochondrial metabolism contradicts the expected outcome of mTOR overactivation^8,39^. This suggests that the effect of TSC2 loss-of-function might be more complex than mTOR simply being overactivated, where brain homeostatic processes are likely to modulate the loss-of-function impact. It is likely that homeostatic processes are modulating the mTOR pathway and influence cellular metabolism as we detected a decrease in expression in the nutrient-sensing mTOR pathway members RRAGA and EEF2^63^ as well as in the major, direct mTOR activator RHEB^8^. Therefore, there might be not enough energy to produce amino acids, as the genes of upstream sensor of amino acids *RRAGA* and downstream translation activation *EEF2* are downregulated. This might consequently lead to protein biogenesis becoming greatly downregulated across all neuronal subtypes in TSC. Concomitantly, homeostatic processes counteract mTOR hyperactivation and decrease *RHEB* gene expression.

It was peculiar to see the largest effect of the metabolic switch in principal neurons, as well as in PVALB and VIP GABAergic neurons, since we previously found similar neuron types to be the most affected in another focal epilepsy - the dysplastic cortex associated with TLE^22^. Moreover, GWAS genes linked to focal, but not generalized, epilepsy were also enriched in PVALB and VIP subtypes. Thus, increased excitation of principal neurons, with reduced inhibition of PVALB neurons and enhanced disinhibition by VIP neurons, might constitute a common seizure-generating circuit in cortical focal epilepsies.

The largest magnitude of gene expression changes was found in subtypes of principal neurons and the ID2 family of GABAergic neurons (Fig. 3a). While it was expected for principal neurons, the discovery that ID2 family neurons, primarily residing in layer 1^64^, are associated with epilepsy, was unexpected and novel, as they have not previously been reported play a role in epileptogenesis. Interestingly, ID2 neurons are the last to mature in the human cortex^65^ and they originate from the caudal ganglionic eminence^64^, which was recently proposed as the major lineage expanded in TSC during development^66^, again indicating particular vulnerability of late cortical development in TSC.

Strikingly, another common feature of both focal epilepsy in TSC (this study) and in TLE^22^ was the strong upregulation of glutamate signaling in principal neurons and, in particular, AMPA receptor auxiliary subunits^67^, CKAMP44 (gene *SHISA9*), CKAMP52 (*SHISA6*), GSG1L (*GSG1L*), TARPg2 (*CACNG2*), and TARPg3 (*CACNG3*). As in TLE cortex, several members among the CKAMP AMPA receptor auxiliary proteins^68^ were upregulated. Thus, CKAMP44 was upregulated across all layers of principal neurons (Fig. 3f). Increases in AMPA receptor-bound CKAMP44 might decrease the deactivation rate for AMPA receptors, increase synaptic strength, and modulate short-term synaptic plasticity^69,70^. The upregulation of another CKAMP family member – CKAMP52 – might increase surface expression of AMPA receptor subunits and increase glutamate potency^68^.

Although all subtypes of GABAergic neurons were preserved, few specific subtypes exhibited changes in overall abundance. Interestingly, we mostly detected an increased abundance in the TSC lesions, which included GABAergic neurons representing PVALB_MEPE and PVALB_LGR5 neurons as well as a rare SST_TH subtype. PVALB_MEPE and PVALB_LGR5 are basket cells^32^ and project onto the somas of excitatory principal neurons, providing a strong inhibitory tone^71^. Intriguingly, these neurons might be particularly sensitive to metabolic impairments in TSC due to high energy demands^72^. SST_TH neurons were previously reported as deep-layer GABAergic neurons that are present in human cortex while absent in great apes^33^. In human cortex, these neurons express TH protein and might release dopamine, whereas SST was detected at the RNA but not at the protein level^33^, which also might be due to very low levels of protein expression^73,74^. Changes both in basket and in SST_TH neurons can lead to functional modulation of local circuits, where the former might be a homeostatic mechanism to increase inhibition within the epileptic focus and the latter might lead to higher levels of dopamine within cortical tubers.

All three major types of glial cells exhibited significant changes in TSC lesions. The increase in microglia observed in the snRNA-seq data confirms previous neuroanatomical reports of elevated microglia numbers in TSC lesions^34,35^; however, we did not detect an upregulation of proinflammatory pathways in TSC microglia. We also demonstrated an increase in a subpopulation of oligodendrocytes in TSC lesions. Interestingly, prior studies reported hypomyelination^75^ and oligodendrocyte loss^76^ in TSC. Our data show that metabolism and protein biogenesis are impaired in TSC oligodendrocytes, thus indicating that a subpopulation of the TSC-elevated oligodendrocytes we identified might represent dysfunctional oligodendrocytes that do not contribute to myelination. Finally, we observed an overall reduction in astrocytes in TSC, which might be either a trait that was overlooked by classical histology methods or could be due to size exclusion during nuclear sorting (see Methods) and disruption of large cytomegalic cells that represent TSC astroglia.

Overall, our study demonstrates that, despite the highly disorganized structure of TSC, all neuronal identities present in the neurotypical cortex are also present in TSC, although the compositional and transcriptomic landscape is altered. This, in turn, opens tremendous potential translational opportunities, whereby local modulation of circuit reorganization can potentially restore circuit structure and ultimately recover function and ameliorate seizures. Additionally, therapeutic opportunities might involve modulating AMPA signaling and, in particular, AMPA auxiliary subunits, as well as offer the perspective of a ketogenic diet that takes into account our findings for downregulation mitochondrial respiration and switching to fatty acid metabolism.

## Acknowledgements

The authors are grateful to the members of the Khodosevich, Aronica, and Petanjek groups for critical input on this project. We wish to thank BRIC Single Cell Genomics (Bente Møller) and Imaging Core Facilities (Yasuku Antoku) for excellent help during this study. This work was supported by the Novo Nordisk Foundation Hallas-Møller Investigator Grants - Emerging NNF16OC0019920 and Ascending NNF21OC0067146 to KK, Lundbeck Foundation Ascending Investigator Grant 2020-1025 to KK, and ZonMw Programme Translational Research no. 95105004 to EA. The work of TLP and ZP was supported by the Croatian Science Foundation Grant No. IP-2022-10-8493

## Methods

### Sample information

The description of human brain samples is provided in Extended Data Table 1.

Cortical tubers were obtained from tuberectomy of frontal lesions of patients with *TSC2* loss-of-function variant tuberous sclerosis undergoing surgery at the Department of Neuropathology at Amsterdam UMC (Amsterdam, The Netherlands), the UMC Utrecht (Utrecht, The Netherlands), and Queensland Children’s Hospital, Australia. In surgery, MRI-evaluated, epileptogenic cortical tubers were resected and immediately flash frozen in liquid nitrogen. Postmortem control tissues were obtained from the archives of the medical centers listed above. The experiments were approved by the Ethical Committee in the Capital Region of Denmark. Tissue was obtained and used in accordance with the Declaration of Helsinki and the Amsterdam UMC Research Code provided by the Medical Ethics Committee and according to the Amsterdam UMC and UMC Utrecht Biobank Regulations (W21-295; TCbio protocol 21-174).

The TSC2 and control samples were selected with respect to several biological covariates such as sex, age, medication, anatomical location, and RNA integrity numbers (RIN). All samples were used for snRNA-seq and selected samples (5 from control and 5 from TSC) were chosen for RNAscope fluorescent *in situ* hybridization due to limited quantities of tissues after determining genotype and performing general pathology for diagnosis. All samples for snRNA-seq were fresh frozen and formalin fixed. For RNAscope, samples were collected from formalin-fixed paraffin-embedded tissue blocks. The anatomical location of resected frontal cortical tubers corresponds to postmortem control samples originating from Brodmann area 10 (BA10).

### Nuclei isolation and FANS

Nuclei were isolated as in^22^. Before fluorescence-activated nuclei sorting (FANS), nuclei were stained with msNeuN-488 antibody (1:1890, Millipore, MAB3777x) for 10 min in darkness at 4 °C. For control of gating, IgG1 κ Isotype (1:188, STEMCELL Technologies, 60070AD.1, 0.2 ug/µL) was used. After staining, nuclei were washed in 0.5% BSA in 1X phosphate-buffered saline (PBS) with RNAse inhibitor (Takara, 2313B, final concentration 0.4 U/μl) and spun down at 1000*g* for 10 min at 4 C°; subsequently, the pellet was resuspended in 100 µL 0.5% BSA/PBS with RNAse inhibitor. Samples were filtered through a 35-µM strainer (Falcon) and 400 µL of 0.5 BSA/PBS with RNAse inhibitor was used to wash the filter to end up with a final volume of 500 µL nuclei suspension. Nuclei were stained with 0.75 µL nucleic acid fluorophore 7-aminoactinomycin (7-AAD) (Sigma, A9400-1MG, 1 mg/mL) into 500 µL of nuclei suspension and incubated on ice until sorting. Prior to sorting, 1.5 mL Lo bind Eppendorf for sorting were coated for at least 24 h with 1.5 mL 0.5% BSA in 1X PBS and aspirated just before initiating the sorting. Sorting was performed on a BD FACSAria III (70-µM nozzle) with cooling set to 4 °C. Nuclei were sorted into two fractions, a NeuN+ for neurons and a NeuN- for glia and other non-neuronal cells. Nucleus suspensions were placed on ice immediately after sorting, nuclei were counted using a hemocytometer, and quality was evaluated. To avoid overrepresentation of oligodendrocytes and thus a corresponding reduction in the number of neurons that are recovered, which is typical for snRNA-seq analyses of the human brain, sorted fractions were pooled together with 20% NeuN- and 80% NeuN+ ratio. A strategy to enrich neurons to 80% of the total cell input was chosen due to limited availability of the rare, resected pediatric brain tissue. In order to identify and characterize the most subtypes of neurons, including rare ones, a relatively high number of neurons is required to gain the resolution and granularity to capture neuronal populations that are low in abundance.

### 10x Chromium library preparation

Chromium Single cell 3’ Reagent Kits v3.1 (10x Genomics, PN-1000121) were used to prepare the library in accordance with the standard protocol. Sorted nuclei were loaded onto the Chromium Next GEM Chip G. On average per sample, 12,419 nuclei were added to individual wells on the Chromium chip to yield on average 8,200 cells after sequencing (Extended Data Table 1). For quantification of cDNA and library concentrations and quality, the Qubit HS dsDNA Assay Kit (Thermo Fisher Scientific, Q32854) and Qubit Fluorometer as well as the High Sensitivity DNA Kit (Agilent, 5067-4626) and Agilent 2100 Bioanalyzer were used. Sample index polymerase chain reaction (PCR) was performed with 12 cycles.

### Sequencing

Libraries were pooled such that the expected amounts of nuclei and expected reads per nuclei matched on the two 100-cycle NovaSeq 6000 S2 flow cells (Illumina, 20012861) for sequencing on an Illumina NovaSeq 6000 (Illumina, 20012850). Cycle setup for sequencing was read1/i7/i5/read2 -> 28/10/10/90. Mean number of reads, unique molecular identifiers (UMI), and genes were 45,944, 11,610, and 3,151 per nucleus, respectively, after *Cellbender* droplet calling.

### Data preprocessing

Quality of sequenced libraries was evaluated via the Illumina base space browser platform and the *Cellranger v7.0.1* mkfastq and count pipelines were used to map reads, demultiplex samples, and call nuclei. Reference transcriptome Human GRCh38 (GENCODE v32/Ensembl98) (refdata-gex-GRCh38-2020-A) provided by 10x Genomics was used for the pipeline and can be downloaded through: wget “https://cf.10xgenomics.com/supp/cell-exp/refdata-gex-GRCh38-2020-A.tar.gz” in unix terminals.

Raw feature barcode matrices were used for further preprocessing with the ambient RNA removal tool *Cellbender v0.3.0*^77^ (https://github.com/broadinstitute/CellBender) provided through *CRMetrics v0.3.1*.^78^ (https://github.com/khodosevichlab/CRMetrics).

Samples were processed using the *remove-background* module. Script can be found in Data Availability.

After assessing and removing ambient RNA counts, a total of 122,731 nuclei were called. Secondary preprocessing was performed using *pagoda2 v1.0.11 (*https://github.com/kharchenkolab/pagoda2) and *Conos v1.5.0* (https://github.com/kharchenkolab/conos).

Subsequent preprocessing was carried out in a cell type-based dividing approach where initial preprocessing was carried out on the whole dataset and then split neurons and glia cells into separate count matrices for further processing of depth filtering. As droplet-based scRNA-seq workflows are prone to produce doublets, these were labeled using *scrublet,* whereby 3,444 possible doublets were detected out of 122,731 nuclei. Nuclei with >5% mitochondrial gene fraction (MGF) were subsequently identified with 2,161 nuclei out of 122,731 exceeding the threshold. Doublets and high-MGF nuclei were filtered, leaving 117,135 nuclei for further preprocessing. The data were subsequently split into neurons (*RBFOX3*, *SLC17A7 and/or GAD1 positive*) and glia (AQP4*, MOG, VCAN,* or *CSF1R* positive, for astrocytes, oligodendrocytes, OPCs, or microglia, respectively) for depth filtering as the difference in transcriptome complexity of the cell types results in different depth profiles. Nuclei were filtered with thresholds of 1700< and 600< UMI per nuclei for neurons and glia, respectively, and sample-wise count matrices were concatenated, resulting in a total of 100,282 nuclei after filtering.

### Differential analysis for control vs TSC conditions

Most differential analyses were performed using *Cacoa v0.4.0* (https://github.com/kharchenkolab/cacoa) package for cross-condition analysis (each type of analysis is described below). The detailed description of the methods for the package are described in^31^. Note that the tools from the package have been implemented originally in^22^, successfully re-implemented by other groups in^79–81^, and further successfully implemented by the developers at full-scale of the toolkit in^17,82,83^.

### Covariate analysis

To assess potential impacts of covariates on the compositional and transcriptomic landscape of samples, we conducted a rigorous covariate analysis of the metadata and technical processing information obtained using *Cacoa v0.4.0* (https://github.com/kharchenkolab/cacoa). Sex, age, designated sequencing flow cell, and 10x Genomics Chromium processing steps one, two, and three were tested against condition. Two workflows CoDA and expression distances were employed for testing using standard pipelines as in *Cacoa*^31^.

### sn-RNA-seq data analysis

To annotate cell types in the dataset, clusters were estimated using the unsupervised Leiden algorithm and annotated with canonical cell-type specific markers reported in previous studies^17,22^.

### Compositional data analysis

We conducted compositional data analyses as described in previous work^17,31,82^ by calculating cell loadings with *Cacoa* using the name test parameter “coda” and n.boot for bootstrapping iterations to 1000. The method uses isometric log-ratio (ILR) transformations applied to cell-type fractions, followed by canonical discriminant analysis (CDA) using the *candisc* package to derive weighted contrasts between cell types in TSC and control samples. The bootstrapping ensures the robustness and statistical significance of the separating coefficients with random cell subsampling. In total, as stated previously, 1000 iterations were performed, with each iteration assessing 1000 randomly sampled cells from both the TSC and control groups. To correct for multiple comparisons, Benjamini-Hochberg correction was used to control for the false discovery rate.

### Cluster-free compositional changes

Differential cell density was estimated using *Cacoa*^31^ as in previous studies^17,22^ with method=“graph” set for the function estimateCellDensity. Graph densities were first calculated for each sample and then the difference in sample densities between conditions was estimated for each datapoint using Wilcoxon test statistics. Lastly, the condition labels were permuted and repeated 400 times to estimate significant changes for each data point. As many data points are produced, normal correction for multiple comparisons cannot be used as it results in a much lower power. Thus, cell graph was used for graph densities and a heat filter for signal smoothing was applied, using a winsorizing level of 1%, and permuted 400 times. Z-scores were calculated as in *Cacoa*^31^ and sample adjusted z-score cell densities are shown in Fig. 2d,e.

### Cluster-based expression shifts

Cluster-based expression distances between cell types across conditions were determined by calculating correlation distances between all sample pairs using pseudo-bulk expression profiles. To normalize these distances between conditions, the average distance within each condition was subtracted, which adjusts for possible random effects. To estimate statistical significance, 1000 random permutations of the condition labels for samples within each cell-type were performed. This step centers the distances using a background distance distribution, generated by randomizing the condition group assignments across the samples. The resulting p-values were adjusted for multiple comparisons using the Benjamini-Hochberg corrections.

### Cluster-free expression shifts

Non-cluster-based expression shifts are estimated from a graph manifold and in our case, we used the permuted UMAP embedding from the preprocessing with *pagoda2* and *Conos*^84^ and expression shifts were calculated in *Cacoa*^31^. Neighborhoods were calculated for individual cells on the graph in a sample-wise manner, where molecular counts within each sample are aggregated generating a “pseudo-bulk” neighborhood count matrix with “number of samples” x “number of genes”. In this approach, a cell is considered its own neighbor. Samples with less than three cells in a given neighborhood of a given cell are omitted. From the neighborhood count matrix of a given cell, pairwise correlation distances between samples are estimated. Permutations of condition labels per sample were carried out on the entire dataset instead of only samples present in the neighborhood of individual cells. For estimation of p-values, smoothing was applied using the median filter with a winsorizing value of 0.025. Furthermore, expression shift magnitudes were additionally smoothed over the graph with the median filter so that the shifts match p-values to a higher degree, as these are estimated on the smoothed values^31^.

### Differentially expressed genes

Differential gene (DE) analysis was performed on normalized pseudobulk gene counts using *Cacoa*^31^ with estimatePerCelllTypeDE(), which implements the Wald test in *DEseq2* with parameter independentFiltering=F^85^. The method collapses sample-wise count matrices into a joint count matrix to then collapse gene counts per cell type into sample-wise pseudobulk gene counts, increasing the statistical power of downstream tests. Next, differentially expressed genes (DEGs) are determined by condition grouping (control vs TSC), run with standard workflow, and corrected for multiple comparisons by Benjamini-Hochberg method. Per cell-type DEGs are used for downstream workflows of GO, GSEA, TF inference, and interaction analysis and CELLECT/CELLEX cell-type vulnerability analysis.

### Gene ontology and gene set enrichment analysis

DEGs were used for performed GO enrichment analysis and Gene Set Enrichment Analysis (GSEA) using a wrapper for *clusterProfiler*^86^ in *Cacoa*^31^ filtering for the top 500 up- and downregulated DEGs. Enrichment was evaluated for all three types of GO pathways; molecular function (MF), cellular component (CC), and biological pathway (BP) with filtering for adjusted p-values <0.05 after testing for multiple comparisons by Benjamini-Hochberg correction.

### Transcription factor inference analysis

TF enrichment was calculated using a regulon database *CollecTRI*^42^ through the R package *decoupleR*^87^. For this analysis, we included only processed human TF regulons. To estimate TF activity inference from differentially expressed pathways in our data, we extracted z-score gene pairs from our previous DE analysis and used the matrix to run the Univariate Linear model (ULM) workflow in *decoupleR*. Scores were normalized and rescaled from --3 to 3.

### Pathway activity inference

To directly infer classical signaling pathways from our data, we used *PROGENy*^49^, a curated pathway to target collection with weighted interactions based on species, through a wrapper in *decoupleR*^87^. Here, we used *PROGENy*’s top 500 responsive genes ranked by p-value. To infer only differentially expressed pathways we used z-scores from DE analysis as input instead of a count matrix to the Multivariate Linear Model (MLM). Scores were normalized and rescaled.

### Genetic risk factor cell type vulnerability analysis

Although patients in our cohort have loss-of-function mutations in the *TSC2* gene and show clear causality with tuberous sclerosis presentation in the clinic, the severity of symptoms varies in many patients. We hypothesized that this could be due to contribution from epileptogenic variants in the patient’s genetic background. In an attempt to elucidate this difference in a cell type-specific manner, we integrated our snRNA-seq data with the largest GWAS study on complex epilepsies^50^ (data:http://www.epigad.org/download/final_sumstats.zip) encompassing 29,944 patients with epilepsy and 52,538 controls with our snRNA-seq data using CELLECT^51^. As the epileptogenic nature of TSC is focal, we included GWAS summary reports from European (Caucasian) cohorts with focal epilepsy and included genetic generalized epilepsies (GGE) from patients of the same ethnic group to serve as inter-disease group control data. First, cell-type expression specificity profiles of TSC2 samples were computed using CELLEX. Next, summary statistics of focal epilepsies were processed using CELLECT. Then, we used the MAGMA^88^ workflow for CELLECT. Here, single nucleotide polymorphisms (SNPs) were mapped onto protein-coding gene locations of human reference genome hg19 build 17 and p-values were calculated based on other gene-level metrics and neighbor gene correlations. Lastly, cell-type prioritization analysis was carried out by fitting a linear regression model between calculated MAGMA ZSTATs calculated in the previous step with the specificity values from the CELLEX cell-type expression specificity profiles for each cell-type annotation. This yields p-values which were then subjected to an α-threshold of significance determined by Bonferroni correction. Cell types with lower p-values implied higher vulnerability to SNPs in gene loci that are important for normal cellular function.

### RNAscope single-molecule fluorescent *in situ* hybridization

To validate layer specific excitatory neuronal markers, five control (CTRL) Brodmann area 10 DLPFC and five TSC2 cortical tuber anatomically matched samples were sourced from the same material as included in the snRNA-seq study. Selected samples were matched for variables such as sex and age. Archival sample material had been formalin fixed, paraffin embedded, and cut to 10 µM of width shortly after initial resection.

Three markers were chosen for identifying layer-specific excitatory neurons: *CUX2* for layer 2-3, *RORB* for layer 4, and *FEZF2* for layer 5-6. The probes were paired for triple target identification using the RNAscope multiplex fluorescent reagent kit v2 (Cat. #323100, ACDbio). The following steps were carried out: samples were first fixed in 4% paraformaldehyde in 1xPBS at 4°C for 15 min, then washed twice in 1X PBS on a rocker. The slides were gradually dehydrated in ethanol, starting with a 50% v/v wash for 5 min, followed by 70% v/v, and ending with two washes in 100% v/v ethanol for the same amount of time. After drying on absorbent paper for 5 min, a hydrophobic barrier was drawn around the tissue sections and allowed to dry for another 5 min. Next, the slides were incubated in hydrogen peroxide for 10 min at room temperature (RT), and then rinsed twice with 200 mL of DEPC-treated MilliQ water (Cat. #ICNA0215090225, WVR).

Following this process, the tissue slides were treated with RNAscope Protease IV for 30 min at RT and rinsed twice with 200 mL of 1X PBS. The probes for *CUX2* (C1), *RORB* (C2), and *FEZF2* (C3) were prewarmed at 40°C for 10 min and mixed at a 1:50 ratio. A 1X wash buffer was prepared by warming two 60-mL bottles of 50X wash buffer at 40°C for 10-20 min and diluting them into 6 L with RNase-free water. The C1 and C2 probes were hybridized serially by incubating the slides at 40°C for 2 h, followed by two washes in 200 mL of 1X wash buffer.

The amplification reagents (Multiplex FL v2 AMP 1, AMP 2 and AMP3) were hybridized by incubating the slides at 40°C for 30 min, followed by two additional washes. Fluorescent signals were developed using horseradish peroxidase (HRP) (RNAscope Multiplex FL v2 HRP-C1 for channel 1, HRP-C2 for channel 2, and HRP-C3 for channel 3), incubated at 40°C for 15 min, and washed again with 1X wash buffer. Opal dyes 1:1500 in Tyramide signal amplification (TSA) buffer (Opal 520, Cat. #FP1487001KT, Opal 570, Cat. #FP1488001KT and Opal 690, Cat. #FP1497001KT, Akoya Biosciences) were then applied, with Opal 520 corresponding to C1, Opal 570 to C2, and Opal 690 to C3. The slides were incubated at 40°C for 30 min, washed, and HRP activity was blocked with the multiplex FL v2 HRP blocker. The slides were then counterstained with DAPI for 30 s at RT, coverslipped using an antifade mount, and stored overnight in the dark at RT.

Whole slide imaging was performed on Zeiss Axioscan Z.1 at 20x magnification.

### Quantification of Immunoreactive Cells

Quantification of immunoreactive cells was performed using Neurolucida 2020 (MBF Bioscience, Williston, Vermont, USA) and Neurolucida Explorer (MBF Bioscience, Williston, Vermont, USA) software on confocal images of triple-labeled immunofluorescent (Green C1-CUX2, Orange C2-RORB and Violet C3-FEZF2) and dapi stained histological slides. Using Neurolucida software, stained cortical slices from 5 control and 5 TSC were analyzed. The cortex was from the pia to the white matter divided into 7 equal bins, with the dimensions being consistent between all analyzed slices. Within each bin positively stained neurons were marked with a correspondingly colored dot, and double positive neurons were marked with both their correspondingly colored dots. Once all neurons in cortical bins of a single slice were marked, the corresponding data file was analyzed using Neurolucida Explorer software. Exact numbers of dots for each neuron subtype, and dot colocalization for the number of mixed identity neuron subtypes within each individual cortical bin was calculated. The results of the quantification were exported as data spreadsheets and compared to determine differences between control and TSC samples.

## Data and code availability

Single-cell RNA-seq data will be deposited at GEO and will be publicly available as of the date of publication. The original code will be deposited at GitHub and will be available for the reviewers during reviewer process.

Any additional information required to reanalyze the data reported in this paper is available from the lead contact upon request.

