## Extended Data Information for "Single-cell profiling of cortical tubers in tuberous sclerosis complex shows molecular structure preservation and massive reorganization of metabolism"

### Extended Data Figures

**Extended Data Fig. 1. Overview of snRNA-seq dataset before ambient RNA removal with CellBender.**
**Extended Data Fig. 2. Unsupervised removal of ambient RNA with Cellbender.**
**Extended Data Fig. 3. Preprocessing of Cellbender-processed samples.**
**Extended Data Fig. 4. Preprocessing of CellBender-processed and doublet- and mitochondrial gene fraction-filtered glia.**
**Extended Data Fig. 5. Preprocessing of CellBender-processed and doublet and mitochondrial gene fraction-filtered neurons.**
**Extended Data Fig. 6. Analysis of major covariate impact on gene expression and cellular composition in snRNA-seq data.**
**Extended Data Fig. 7. Dot-plot representing expression of marker genes for Neu_dev_THY1 cluster showing excitatory but not GABAergic inhibitory lineage markers.**
**Extended Data Fig. 8. Volcano plots for DE genes for each subtype of principal neurons in TSC.**
**Extended Data Fig. 9. Volcano plots for DE genes for each subtype of GABAergic neurons in TSC. Statistics and visualization are the same as Extended Data Fig. 8.**
**Extended Data Fig. 10. Dysregulation of gene expression-coding proteins involved in mitochondrial oxidative phosphorylation.**
**Extended Data Fig. 11. Dot plot representing expression levels of DE genes from GO term "Glutamatergic receptor signaling" in TSC neuronal subtypes.**
**Extended Data Fig. 12. Violin plot showing changes in expression levels of *CXCL14* gene across subtypes of VIP neuronal family.**
**Extended Data Fig. 13. Volcano plots for DE genes involved in axon development in TSC.**
**Extended Data Fig. 14. Heatmap showing DE of genes from mTOR pathway in TSC neuronal subtypes.**
**Extended Data Fig. 15. Heatmap showing enriched transcription factor (TF) networks.**
**Extended Data Fig. 16. Heatmap showing differential interaction strength between neuronal cell types.**

### Extended Data Tables

**Extended Data Table 1. Patient metadata and summary statistics for single nucleus transcriptomics.**
**Extended Data Table 2. Lack of significant differential expression of GO terms related to inflammation in TSC microglia.**
**Extended Data Table 3. DE genes in TSC cell subtypes.**
**Extended Data Table 4. Massive differential expression of GO terms related to mitochondrial electron transport for each neuronal subtype in TSC.**
**Extended Data Table 5. Minor differential expression of GO terms related to glycolysis for each neuronal subtype in TSC.**
**Extended Data Table 6. Minor differential expression of GO terms related to neuronal migration for each neuronal subtype in TSC.**
**Extended Data Table 7. TF-enriched networks in DE genes in TSC neuronal subtypes.**

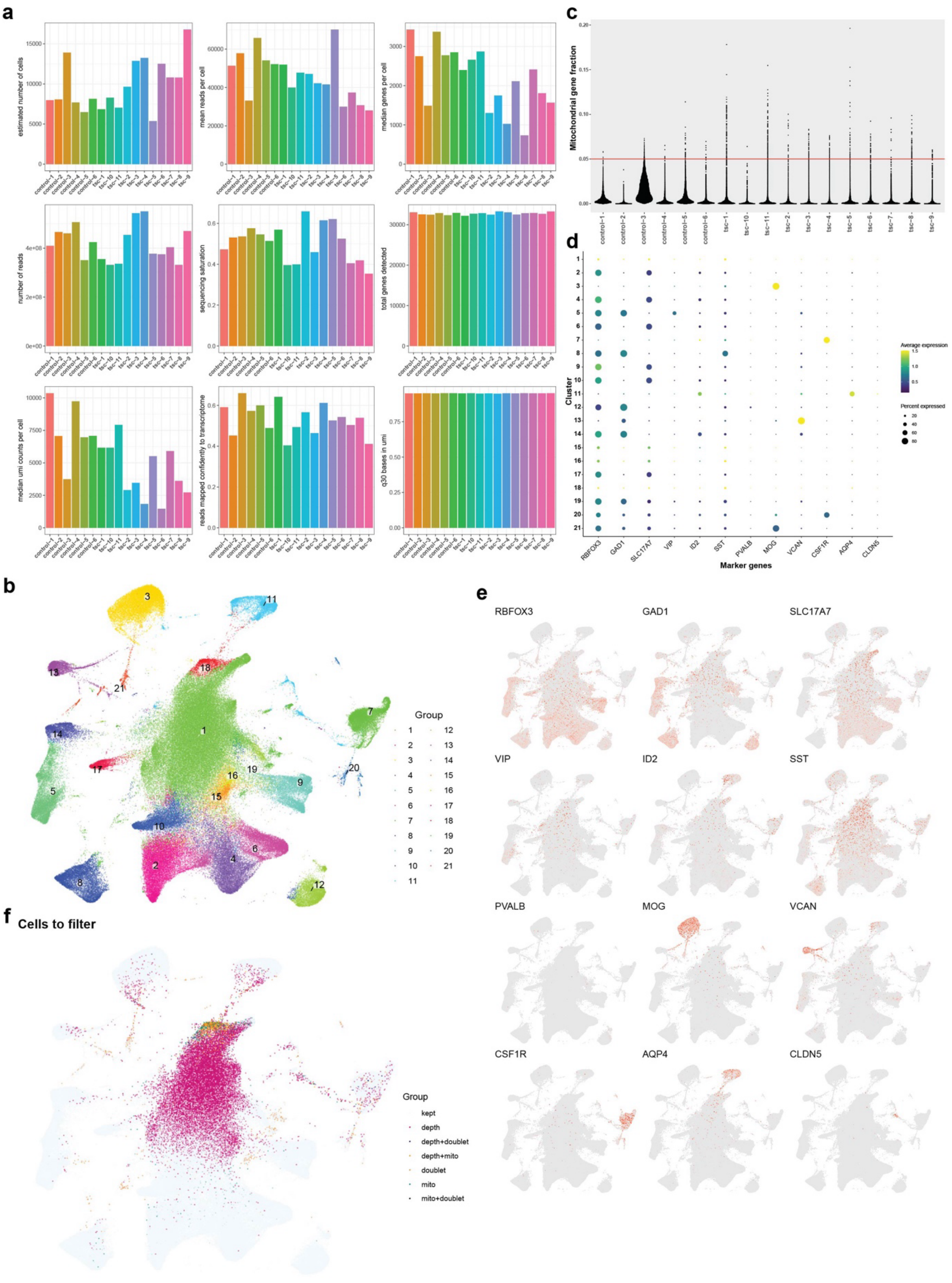

**Extended Data Fig. 1. Overview of snRNA-seq dataset before ambient RNA removal with CellBender. (a)** Sequencing metrics from control (n=6) and TSC (n=11) samples. From top left to right bottom per sample; estimated number of cells, mean reads per cell, median genes per cell, number of reads, sequencing saturation, total genes detected, median UMI counts per cell, reads mapped confidently to transcriptome, and fraction of q30 bases in UMI's.

**(b)** UMAP embedding of filtered count matrix with unsupervised Leiden clustering, resulting in 21 distinct clusters.

**(c)** Scatterplot of mitochondrial gene fraction in cells with a threshold of 0.05 (5%).

**(d)** Dot plot of Leiden clusters and canonical cell-type markers showing average expression and percent of cluster-expressing marker. *RBFOX3* = Neurons, *GAD1* = Inhibitory neurons, *SLC17A7* = Principal neurons, *VIP* = Vasoactive intestinal peptide inhibitory neurons, *ID2* = *ID2* inhibitory neurons, *SST* = Somatostatin inhibitory neurons, *PVALB* = Parvalbumin inhibitory neurons, *MOG* = Oligodendrocytes, *VCAN* = Oligodendrocyte precursor cells (OPCs), *CSF1R* = Microglia, *AQP4* = Astrocytes, and *CLDN5* = Endothelial cells.

**(e)** UMAP embeddings with marker genes of canonical cell types. *RBFOX3* = Neurons, *GAD1* = Inhibitory neurons, *SLC17A7* = Principal neurons, *VIP* = Vasoactive intestinal peptide inhibitory neurons, *ID2* = *ID2* inhibitory neurons, *SST* = Somatostatin inhibitory neurons, *PVALB* = Parvalbumin inhibitory neurons, *MOG* = Oligodendrocytes, *VCAN* = Oligodendrocyte precursor cells (OPCs), *CSF1R* = Microglia, *AQP4* = Astrocytes, and *CLDN5* = Endothelial cells.

**(f)** UMAP embedding of cells to be filtered using standard filtering workflow. Filtering for cells >5% mitochondrial gene fraction, cells classified as doublets, and cells with depth <750 UMIs. This plot illustrates the state of the dataset before removing ambient RNA with CellBender.

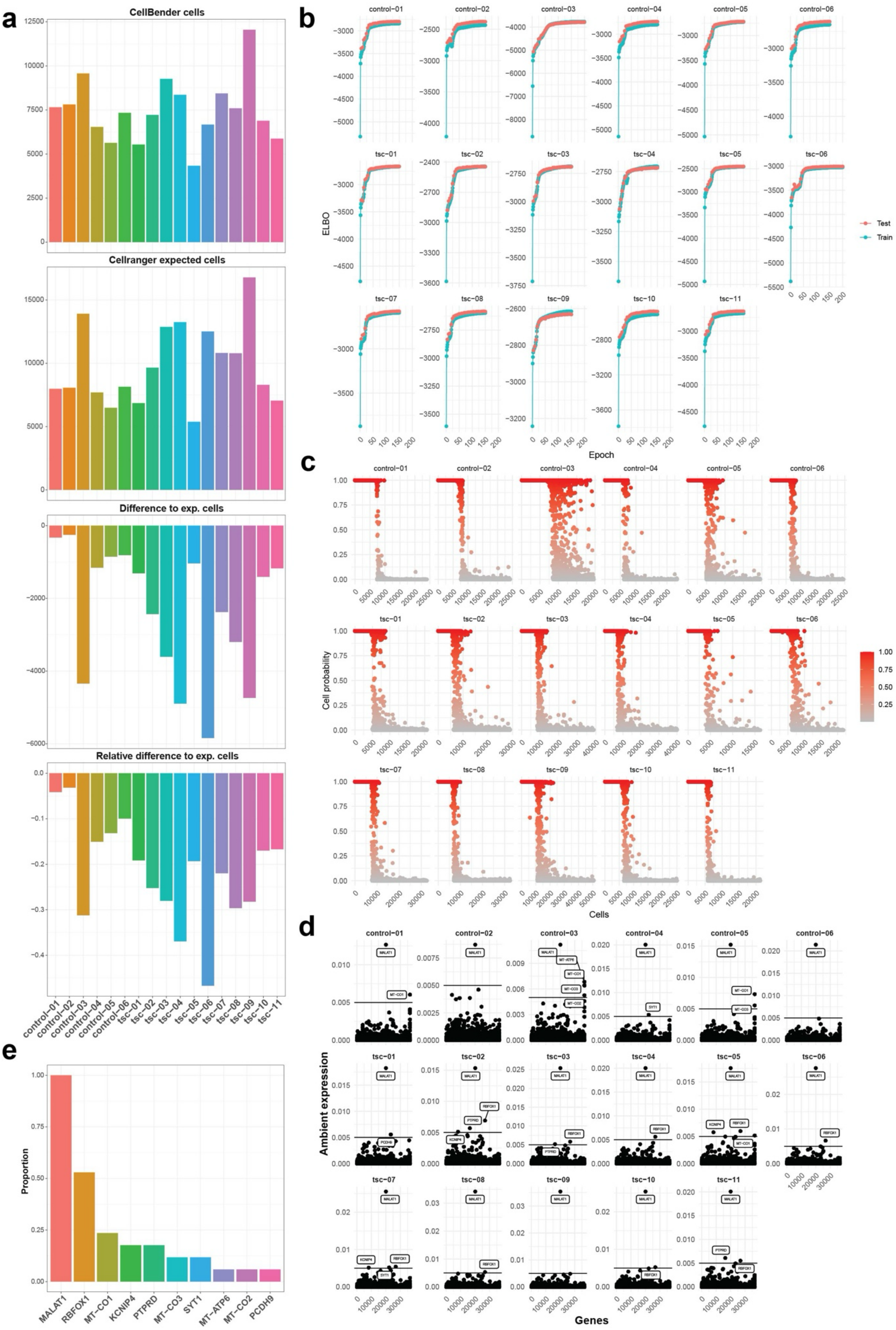


**Extended Data Fig. 2. Unsupervised removal of ambient RNA with Cellbender.**

**(a)** Difference in cell counts per sample (control n=6 and TSC n=11) before and after CellBender processing.

**(b)** CellBender ELBO (test and train) plots ranging from 150 to 200 epochs with varying learning-rates.

**(c)** CellBender cell calling curves displaying cell probabilities of droplets.

**(d)** CellBender sample- wise expression of ambient RNAs.

**(e)** Proportion of samples filtered for specific ambient RNA signatures.

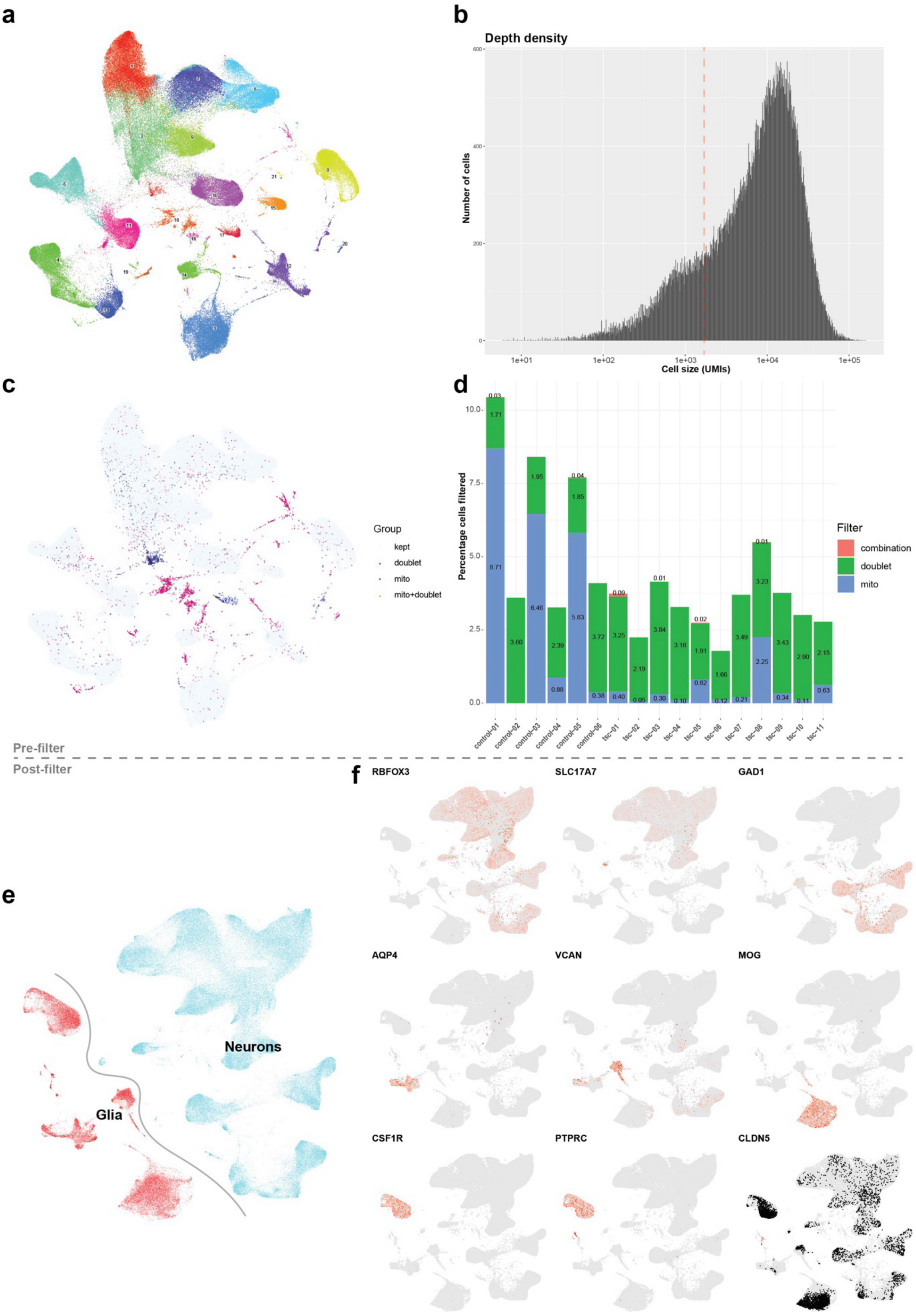


**Extended Data Fig. 3. Preprocessing of Cellbender-processed samples.**

**(a)** UMAP embedding of Cellbender-filtered count matrix with Leiden clustering.

**(b)** Depth bar plot of the entire dataset (control n=6 and TSC n=11). Dotted line at 1700 UMIs indicates division of two cell populations with differing transcriptomic density and complexity. No cells are filtered based on depth at this step.

**(c)** UMAP embedding of cells filtered from doublet calling and/or exceeding >5% mitochondrial gene fraction.

**(d)** Stacked bar plot showing percentage sample-wise filtering from doublet calling, >5% mitochondrial gene fraction, or both.

**(e)** UMAP embedding of postdoublet and mitochondrial gene fraction filtering showing glia in red and neurons in blue. The curved line qualitatively divides the two populations.

**(f)** UMAP embeddings after CellBender processing and doublet and mitochondrial gene fraction filtering with marker genes of canonical cell types. *RBFOX3* = Neurons, *SLC17A7* = Principal neurons, *GAD1* = Inhibitory neurons, *AQP4* = Astrocytes, *VCAN* = Oligodendrocyte precursor cells (OPCs), *MOG* = Oligodendrocytes, *CSF1R* = Microglia, *PTPRC* = Peripheral immune cells (monocytes etc), *AQP4* = Astrocytes and *CLDN5* = Endothelial cells.

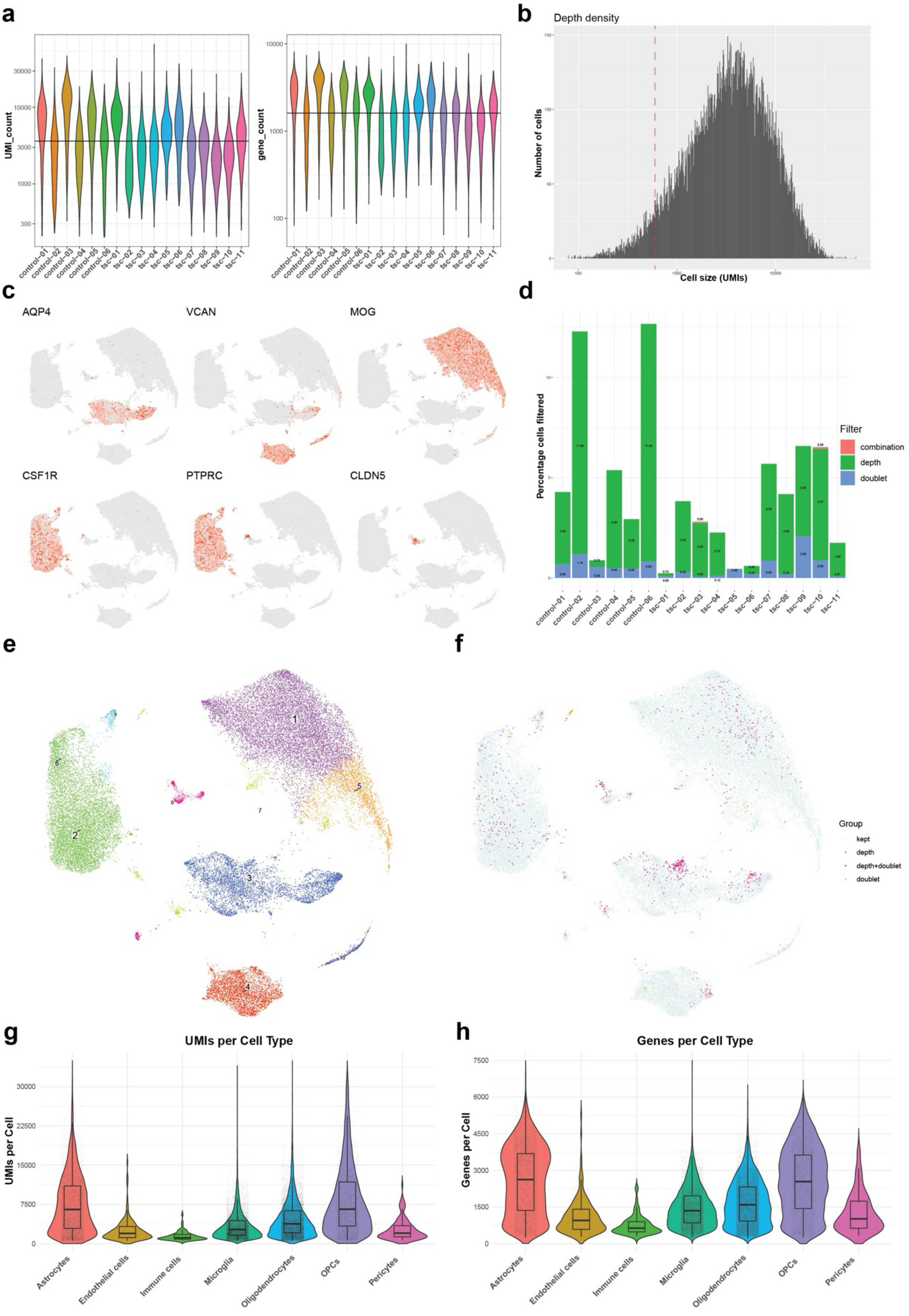


**Extended Data Fig. 4. Preprocessing of CellBender-processed and doublet- and mitochondrial gene fraction-filtered glia.**

**(a)** Violin plots of UMI and gene count per sample (control n=6 and TSC n=11) before depth filtering.

**(b)** Depth bar plot of glial cells, depth threshold set to filter cells with <600 UMIs.

**(c)** UMAP embeddings after CellBender processing and doublet and mitochondrial gene fraction filtering of canonical glial markers. *AQP4* = Astrocytes, *VCAN* = Oligodendrocyte precursor cells (OPCs), *MOG* = Oligodendrocytes, *CSF1R* = Microglia, *PTPRC* = Peripheral immune cells (monocytes etc), *AQP4* = Astrocytes and *CLDN5* = Endothelial cells.

**(d)** Stacked bar plot showing percentage sample-wise filtering of glia with depth <600 UMIs and second round of doublet calling or a combination of the two.

**(e)** UMAP embedding of CellBender-processed and doublet- and mitochondrial gene fraction-filtered glia with Leiden clustering before filtering for depth and second round of doublet calling.

**(f)** UMAP embedding of glia to be filtered by doublet calling or fall below depth threshold of <600 UMIs.

**(g)** Violin plot of UMIs per cell after filtering for canonical glial cell types.

**(h)** Violin plot of genes per cell after filtering for canonical glial cell types.

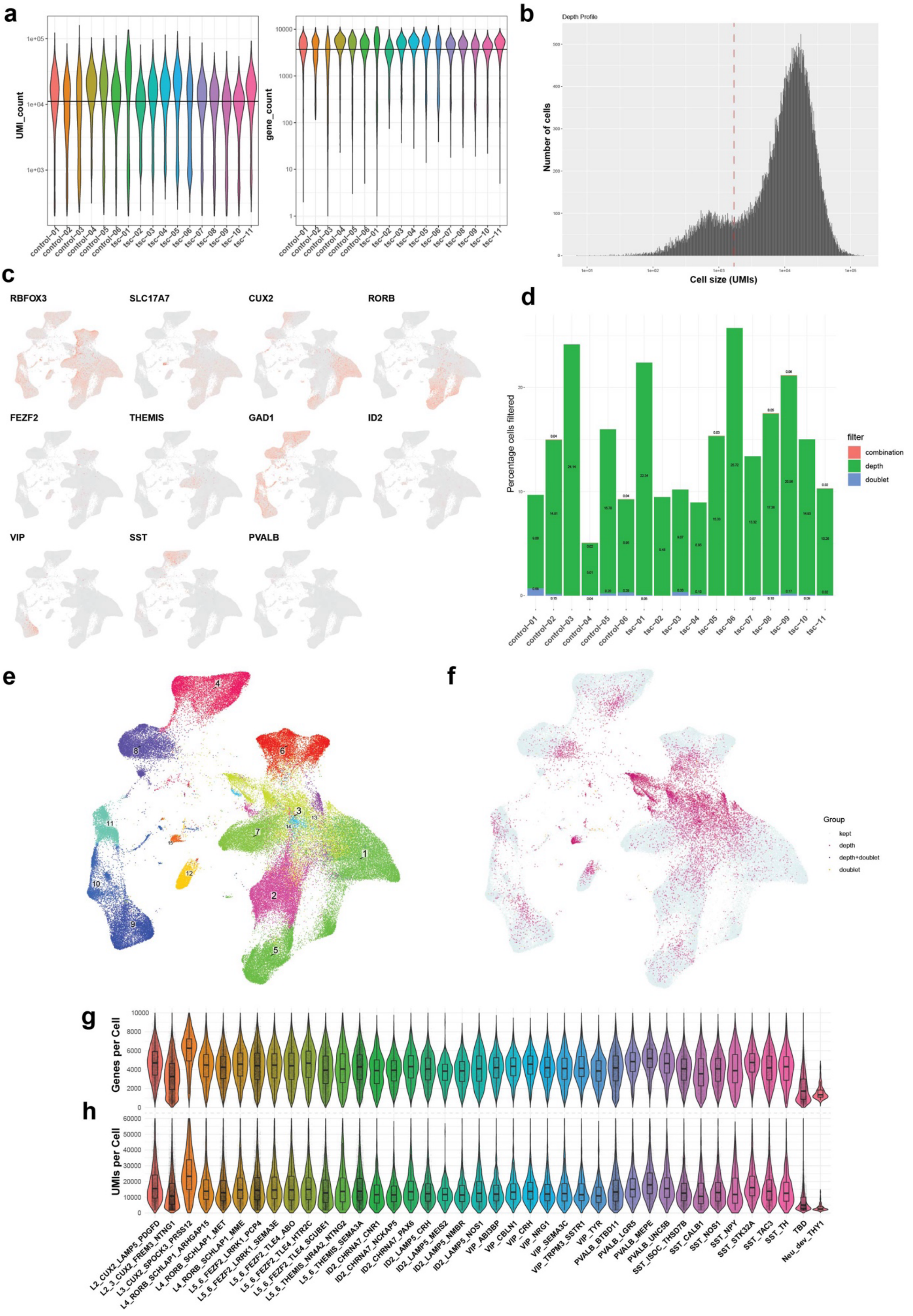


**Extended Data Fig. 5. Preprocessing of CellBender-processed and doublet and mitochondrial gene fraction-filtered neurons.**

**(a)** Violin plots of UMI and gene count per sample (control n=6 and TSC n=11) before depth filtering.

**(b)** Depth bar plot of neurons, depth threshold set to filter cells with <1700 UMIs. The threshold was chosen as a clear separation into two cell populations with differing depth density and transcriptomic complexity was observed at this cell size.

**(c)** UMAP embeddings after CellBender processing and doublet and mitochondrial gene fraction filtering of canonical neuron markers. *RBFOX3* (NeuN) = Neurons, *SLC17A7* = Principal neurons, *CUX2* = Layer 2-3 principal neurons, *RORB* = Layer 4 principal neurons, *FEZF2* = *FEZF2* Layer 5-6 principal neurons, *THEMIS* = *THEMIS* Layer 5-6 principal neurons, GAD1 = Inhibitory neurons, *ID2* = *ID2* inhibitory neurons, *VIP* = Vasoactive intestinal peptide inhibitory neurons, *SST* = Somatostatin inhibitory neurons, *PVALB* = Parvalbumin inhibitory neurons.

**(d)** Stacked bar plot showing percentage sample-wise filtering of neurons with depth <1700 UMIs and second round of doublet calling or a combination of the two.

**(e)** UMAP embedding of CellBender-processed and doublet and mitochondrial gene fraction-filtered neurons with Leiden clustering before filtering for depth and second round of doublet calling.

**(f)** UMAP embedding of neurons to be filtered by doublet calling of fall below depth threshold of <1700 UMIs.

**(g)** Violin plot of UMIs per cell after filtering for high-resolution neuron cell types.

**(h)** Violin plot of genes per cell after filtering for high-resolution neuron cell types.

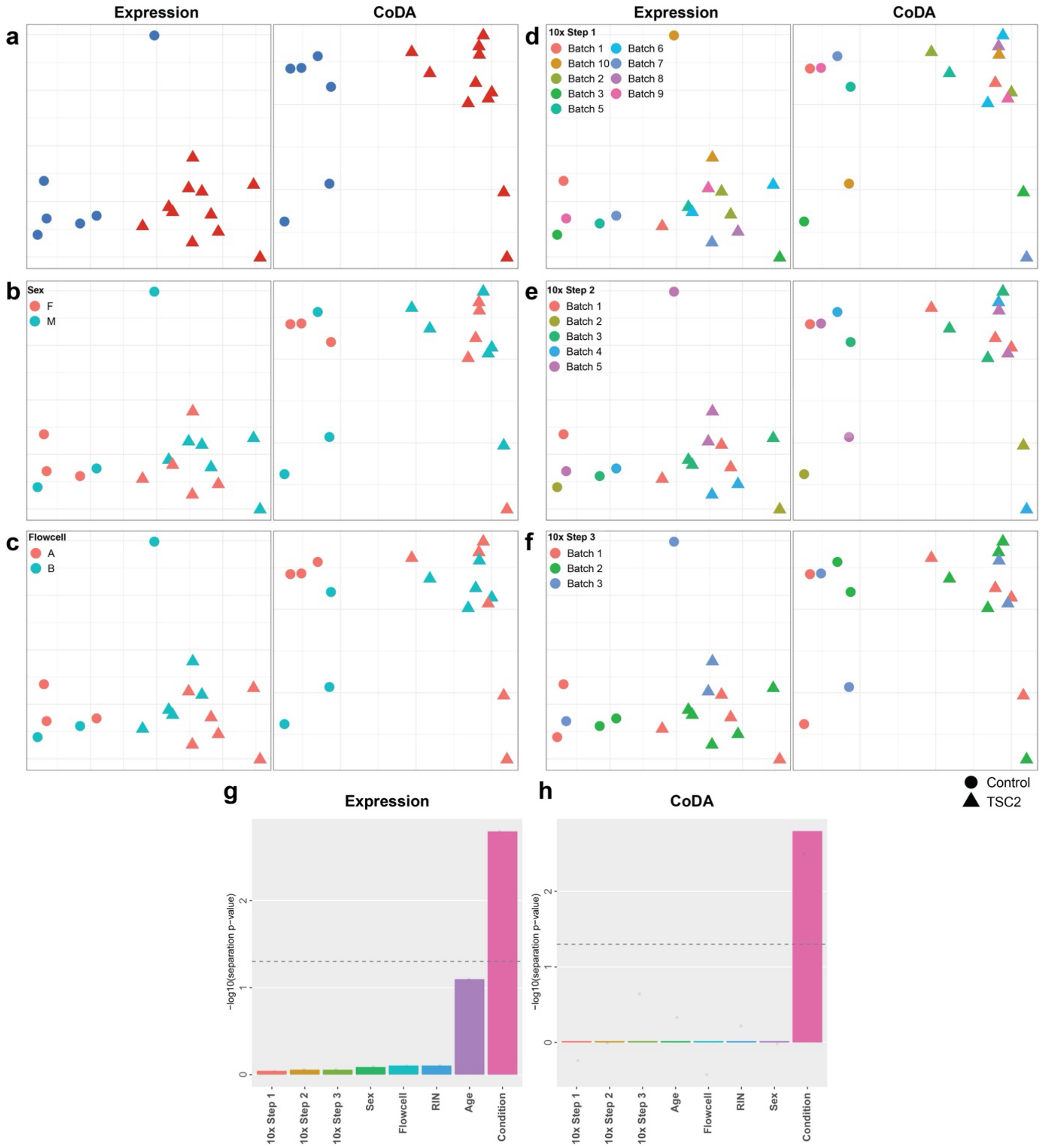


**Extended Data Fig. 6. Analysis of major covariate impact on gene expression and cellular composition in snRNA-seq data.**

**(a)** Multidimensional scaling (MDS) plot visualizing similarity in gene expression (left) and cellular composition by compositional data analysis (CoDA, right) between all samples.

**(b)-(f)** MDS plots are colored by sex (b), sequencing flowcell (c), 10x Genomics protocol step 1: GEM Generation & Barcoding (d), step 2: Post-GEM-RT Cleanup & cDNA amplification (e), step 3: 3' Gene Expression Library Construction (f).

**(g), (h)** Quantifications showing that both gene expression and cellular composition are different across conditions, but not other covariates.

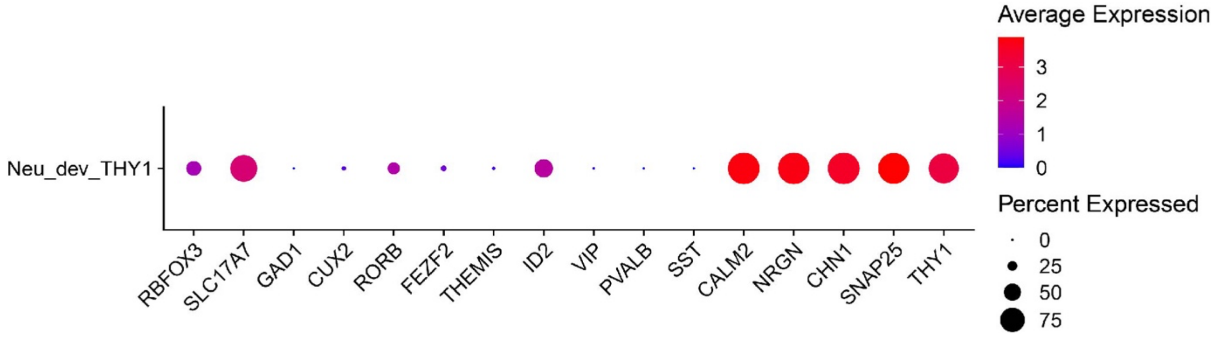


**Extended Data Fig. 7. Dot-plot representing expression of marker genes for Neu_dev_THY1 cluster showing excitatory but not GABAergic inhibitory lineage markers.** Dot size indicates the percent of nuclei that express the marker, and the color represents normalized average expression levels across all Neu_dev_THY1 cells.

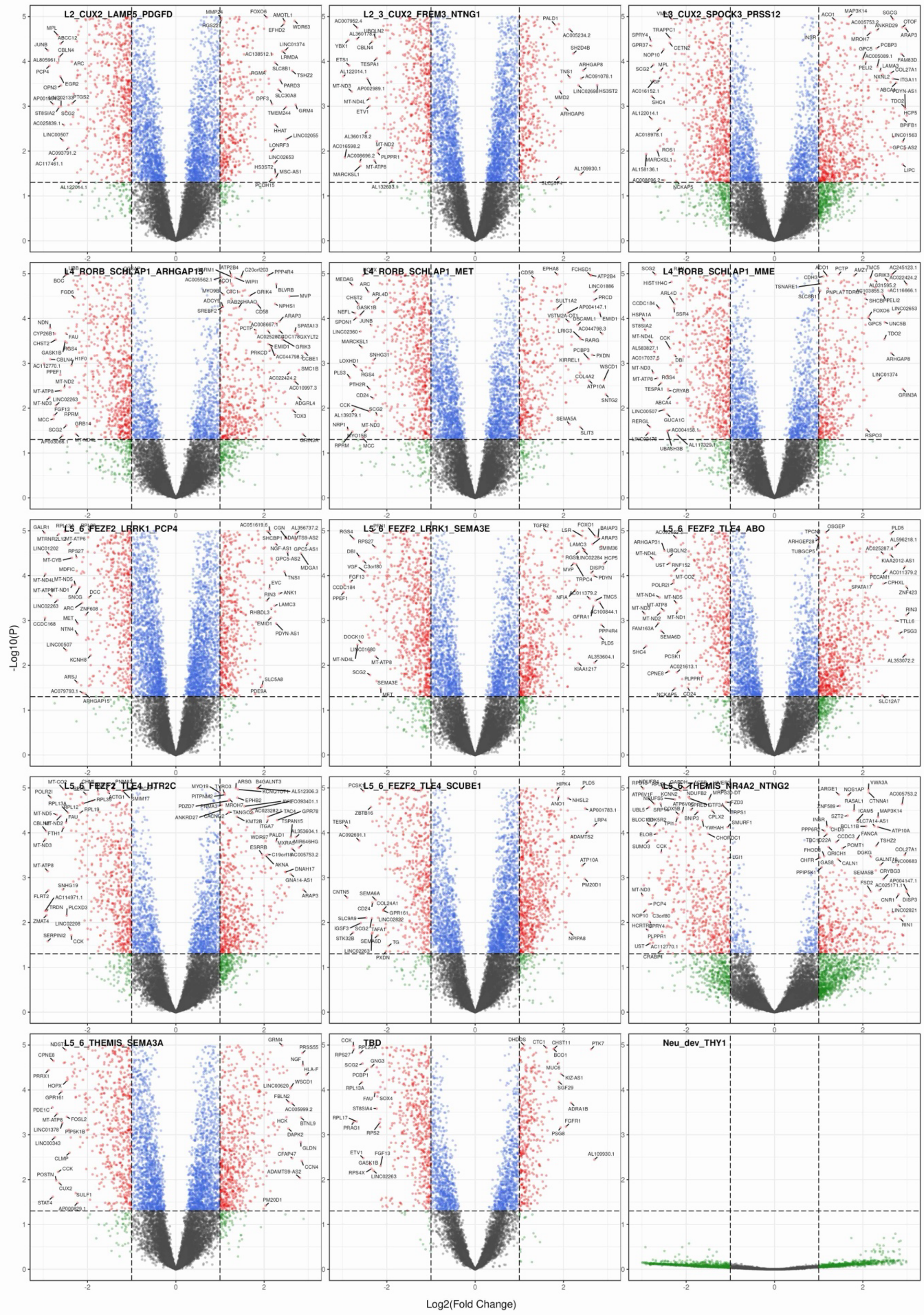

**Extended Data Fig. 8. Volcano plots for DE genes for each subtype of principal neurons in TSC.**

Each dot represents a gene identified using DE analysis. The x-axis is $log_2$-fold magnitude change for gene expression (dotted lines show cut-off $log_2$-fold change of –1.5 and 1.5) and the y-axis is $log_{10}$-fold change for p values (1.3 is ~0.05 p value cut-off). The colors of the dots represent gene expression cell fractions scaled from 0-1. Upregulated and downregulated genes are with positive and negative scale along the x-axis, respectively.

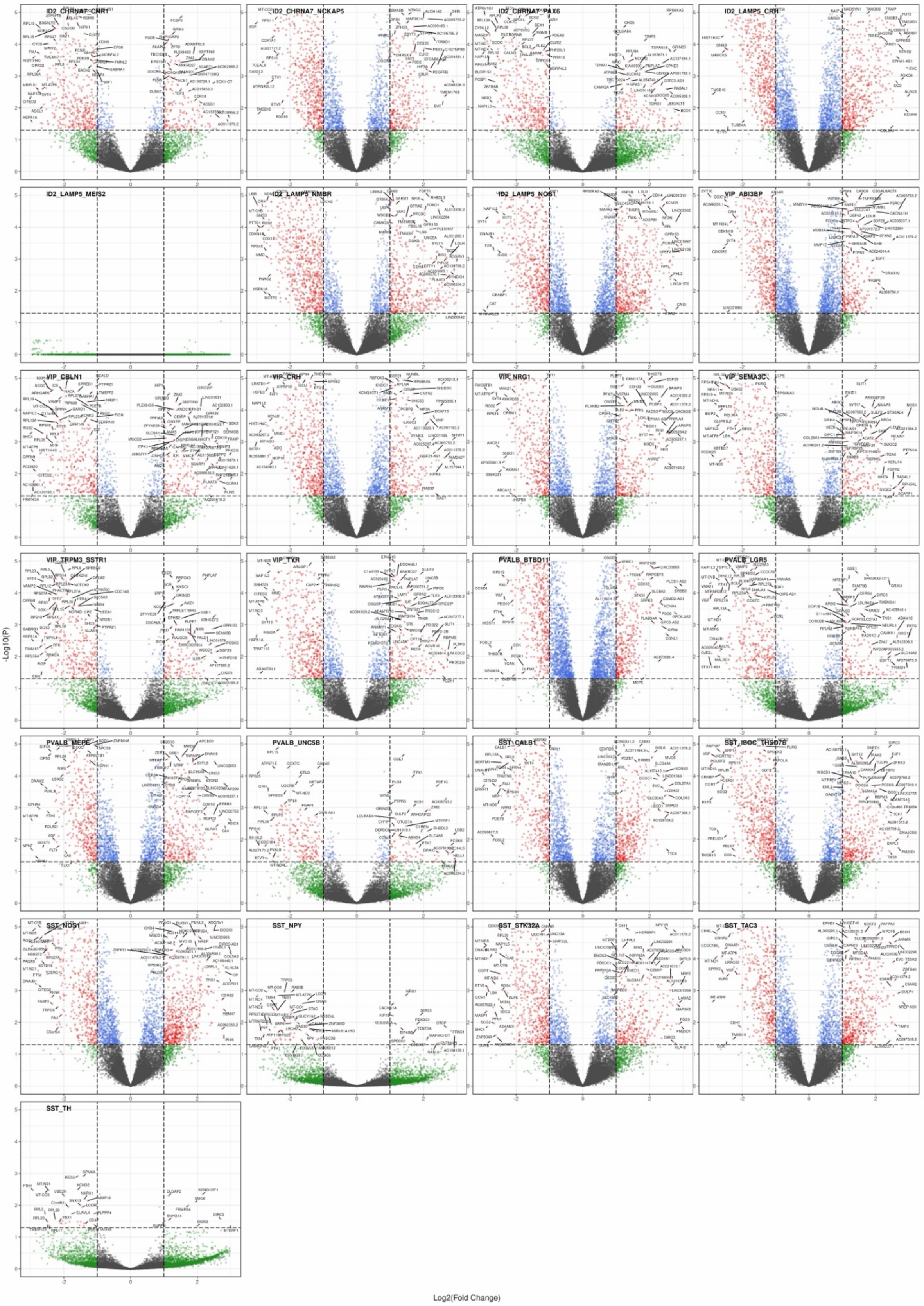


**Extended Data Fig. 9. Volcano plots for DE genes for each subtype of GABAergic neurons in TSC.**

Statistics and visualization are the same as Extended Data Fig. 8.

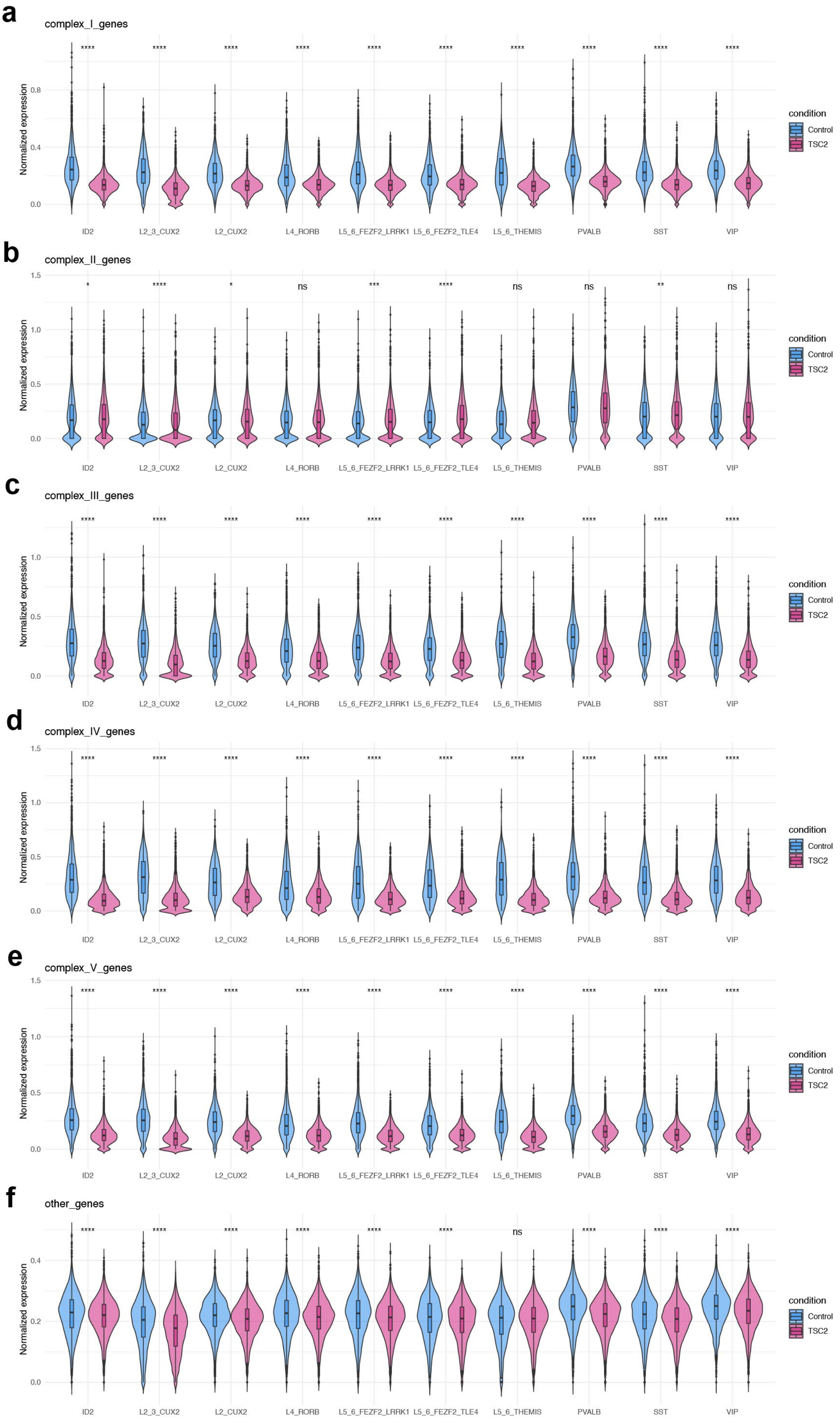
a
complex_I_genes
Normalized expression
0.0
0.4
0.8
**** **** **** **** **** **** **** **** **** ****
ID2 L2_3_CUX2 L2_CUX2 L4_RORB L5_6_FEZF2_LRRK1 L5_6_FEZF2_TLE4 L5_6_THEMIS PVALB SST VIP
condition
Control
TSC2
b
complex_II_genes
Normalized expression
0.0
0.5
1.0
1.5
* **** * ns *** **** ns ns ** ns
ID2 L2_3_CUX2 L2_CUX2 L4_RORB L5_6_FEZF2_LRRK1 L5_6_FEZF2_TLE4 L5_6_THEMIS PVALB SST VIP
condition
Control
TSC2
c
complex_III_genes
Normalized expression
0.0
0.5
1.0
**** **** **** **** **** **** **** **** **** ****
ID2 L2_3_CUX2 L2_CUX2 L4_RORB L5_6_FEZF2_LRRK1 L5_6_FEZF2_TLE4 L5_6_THEMIS PVALB SST VIP
condition
Control
TSC2
d
complex_IV_genes
Normalized expression
0.0
0.5
1.0
1.5
**** **** **** **** **** **** **** **** **** ****
ID2 L2_3_CUX2 L2_CUX2 L4_RORB L5_6_FEZF2_LRRK1 L5_6_FEZF2_TLE4 L5_6_THEMIS PVALB SST VIP
condition
Control
TSC2
e
complex_V_genes
Normalized expression
0.0
0.5
1.0
1.5
**** **** **** **** **** **** **** **** **** ****
ID2 L2_3_CUX2 L2_CUX2 L4_RORB L5_6_FEZF2_LRRK1 L5_6_FEZF2_TLE4 L5_6_THEMIS PVALB SST VIP
condition
Control
TSC2
f
other_genes
Normalized expression
0.0
0.2
0.4
**** **** **** **** **** **** ns **** **** ****
ID2 L2_3_CUX2 L2_CUX2 L4_RORB L5_6_FEZF2_LRRK1 L5_6_FEZF2_TLE4 L5_6_THEMIS PVALB SST VIP
condition
Control
TSC2

**Extended Data Fig. 10. Dysregulation of gene expression-coding proteins involved in mitochondrial oxidative phosphorylation.**

(a) - (f) Violin plots showing DE of complex I (a), complex II (b), complex III (c), complex IV (d), complex V (e), and other genes that are not part of complex I-V (f) between control and TSC cells for each family of neurons. Bars show median with standard deviations; dots label the outliers.

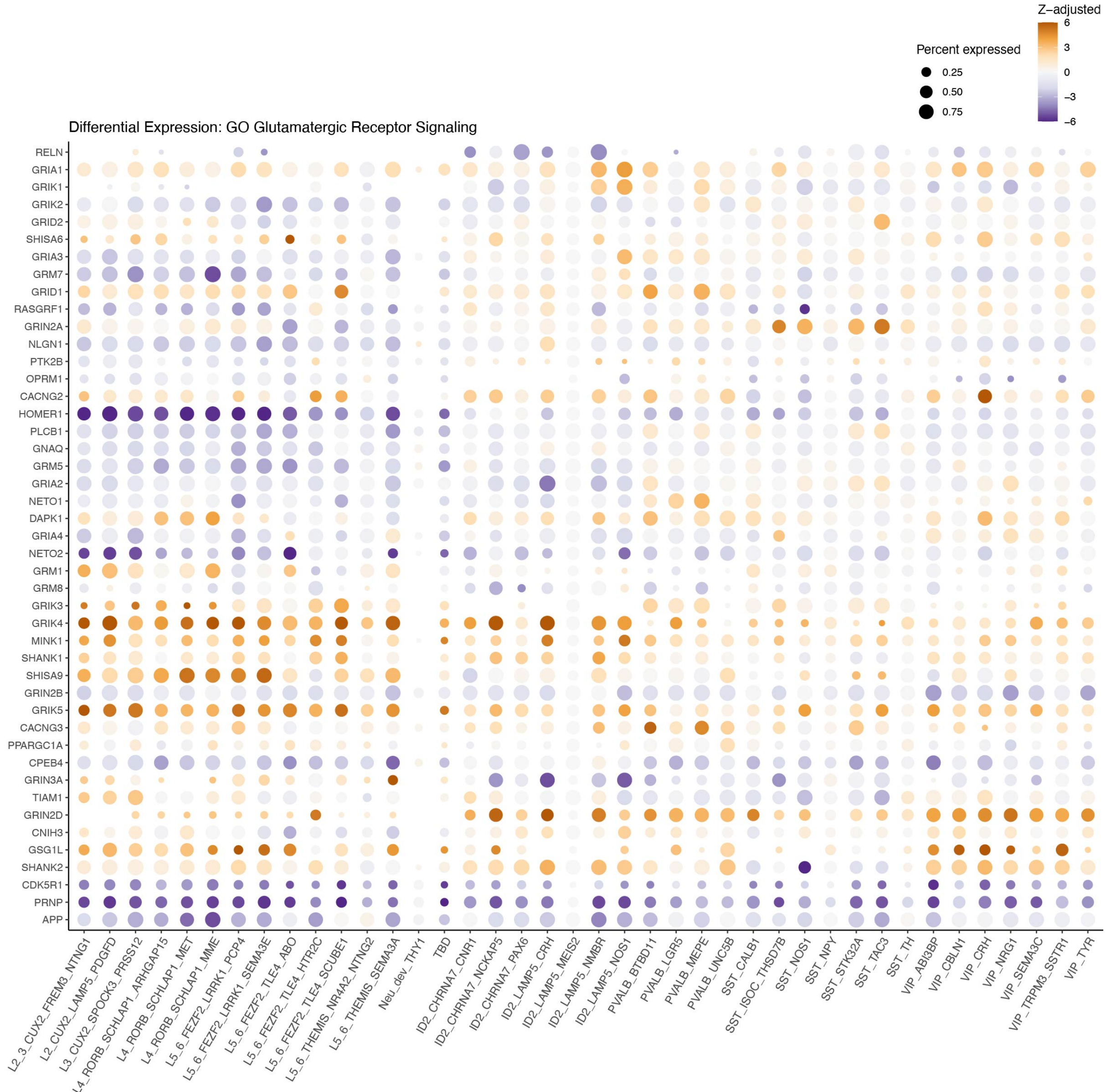


**Extended Data Fig. 11. Dot plot representing expression levels of DE genes from GO term "Glutamatergic receptor signaling" in TSC neuronal subtypes.** Subtypes are ordered by principal neurons followed by GABAergic neurons. The size of the dots reflects the percent of cells within a subtype that expresses a gene. Orange and blue colors indicate up- and downregulation, respectively, and the intensity of the color corresponds to Z-adjusted value for expression change.

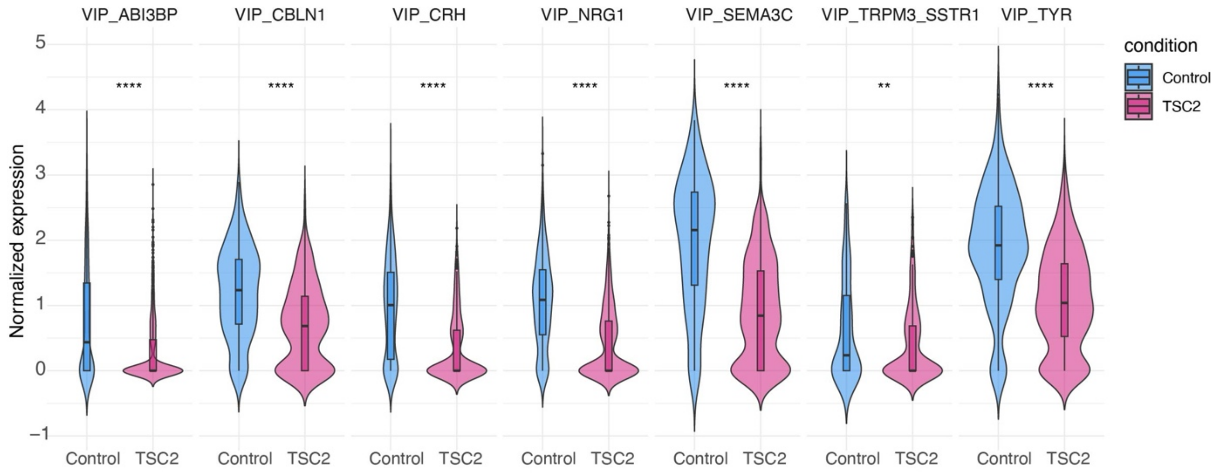


**Extended Data Fig. 12. Violin plot showing changes in expression levels of *CXCL14* gene across subtypes of VIP neuronal family.** Bars show median for absolute $log_2$-fold change values with standard deviations; dots label the outliers.

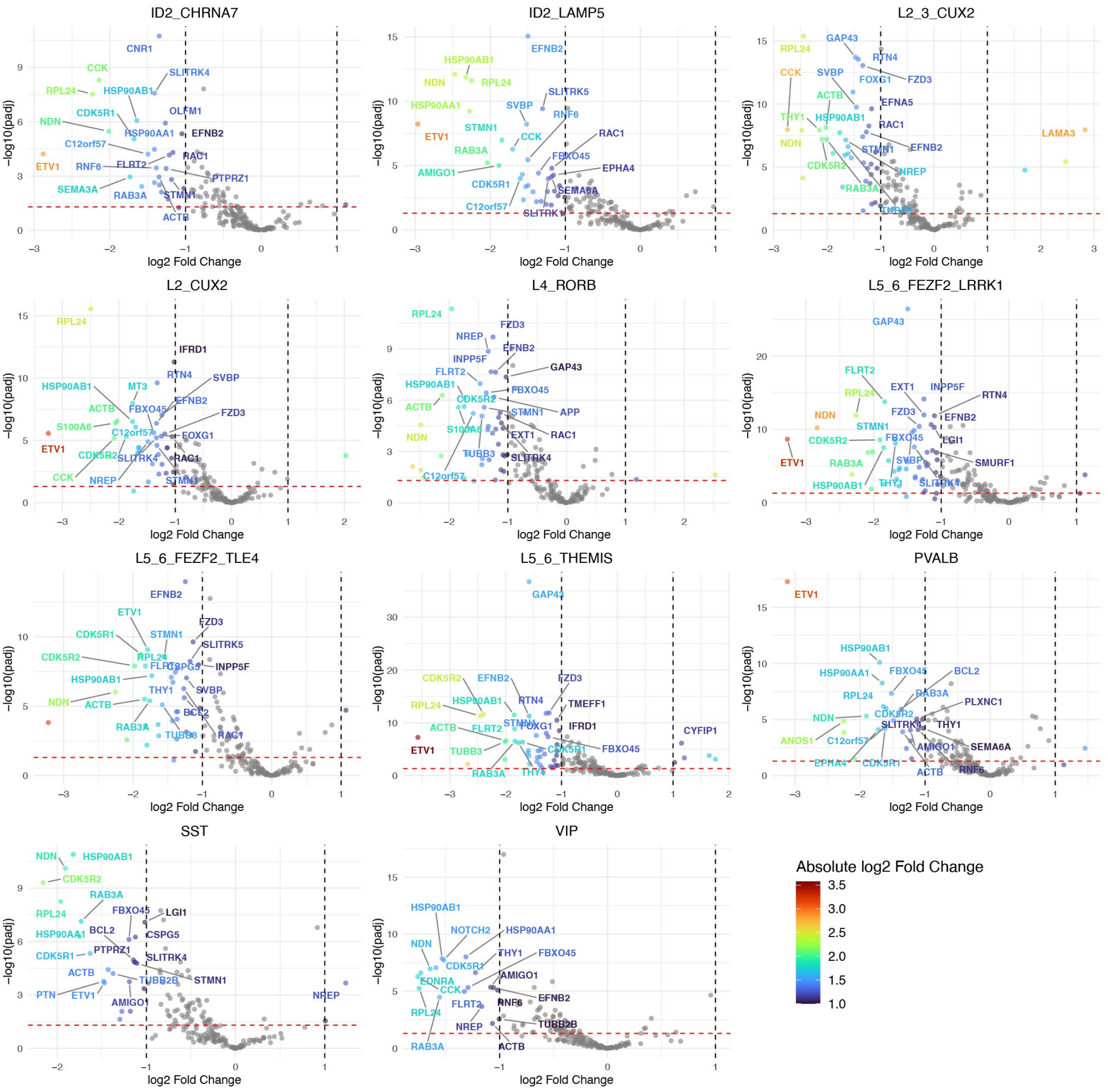


**Extended Data Fig. 13. Volcano plots for DE genes involved in axon development in TSC.**

Each dot represents a gene identified using DE analysis. The x-axis is $\log_2$-fold magnitude change for gene expression (dotted cut-off lines are at 1 and -1) and the y-axis is -$\log_{10}$ -fold change for p values (red line at 1.3 is 0.05 p value cut-off). The colors of the dots represent absolute $\log_2$-fold change values. Upregulated and downregulated genes are with positive and negative scale along the x-axis, respectively. Note that the vast majority of genes are downregulated.

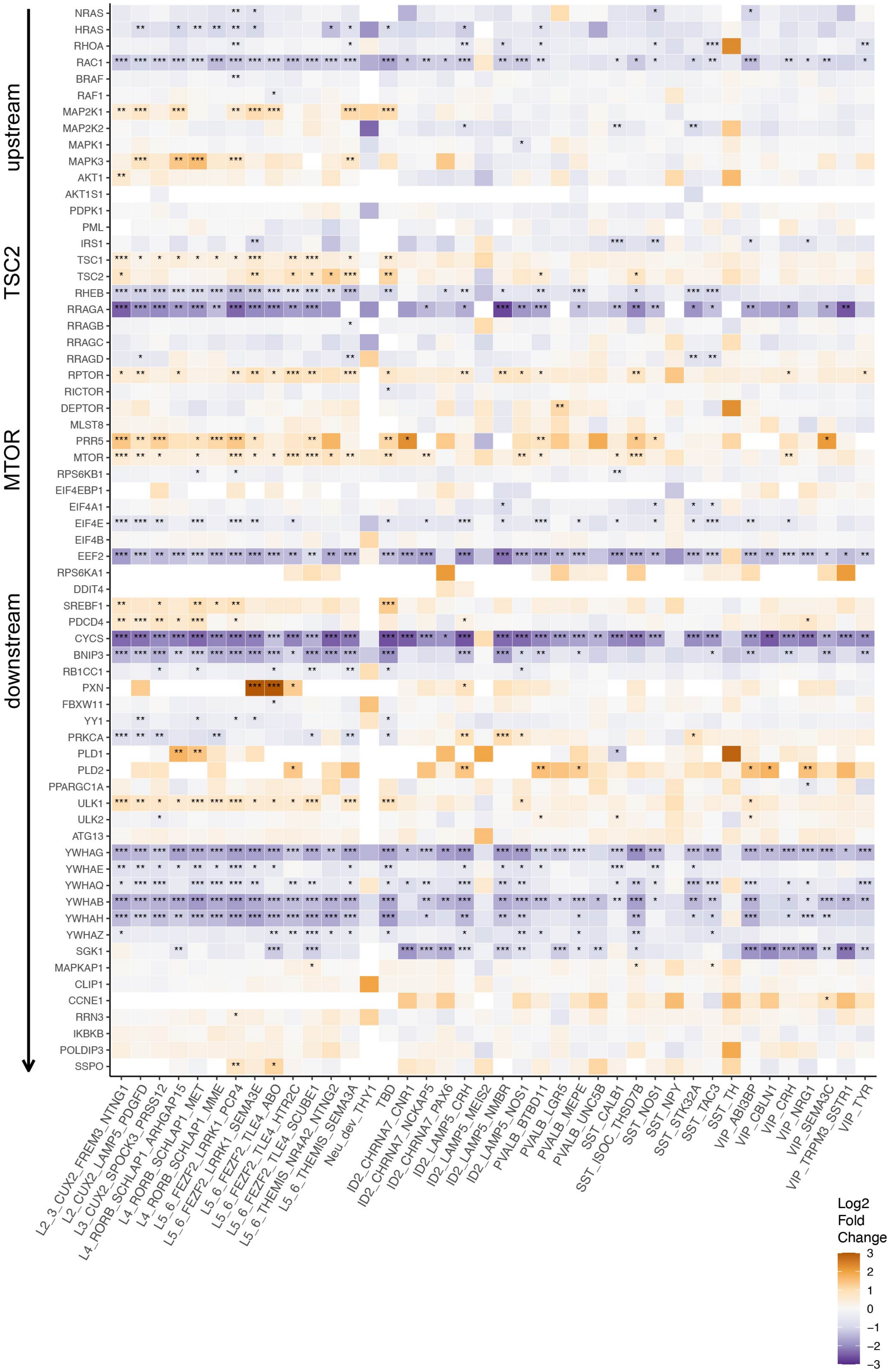
upstream
TSC2
MTOR
downstream
NRAS
HRAS
RHOA
RAC1
BRAF
RAF1
MAP2K1
MAP2K2
MAPK1
MAPK3
AKT1
AKT1S1
PDPK1
PML
IRS1
TSC1
TSC2
RHEB
RRAGA
RRAGB
RRAGC
RRAGD
RPTOR
RICTOR
DEPTOR
MLST8
PRR5
MTOR
RPS6KB1
EIF4EBP1
EIF4A1
EIF4E
EIF4B
EEF2
RPS6KA1
DDIT4
SREBF1
PDCD4
CYCS
BNIP3
RB1CC1
PXN
FBXW11
YY1
PRKCA
PLD1
PLD2
PPARGC1A
ULK1
ULK2
ATG13
YWHAG
YWHAE
YWHAQ
YWHAB
YWHAH
YWHAZ
SGK1
MAPKAP1
CLIP1
CCNE1
RRN3
IKBKB
POLDIP3
SSPO
L2_3_CUX2_FREM3_NTNG1
L2_CUX2_LAMP5_PDGFD
L3_CUX2_SPOCK3_PRSS12
L4_RORB_SCHLAP1_ARHGAP15
L4_RORB_SCHLAP1_MET
L4_RORB_SCHLAP1_MME
L5_6_FEZF2_LRRK1_PCP4
L5_6_FEZF2_LRRK1_SEMA3E
L5_6_FEZF2_TLE4_ABO
L5_6_FEZF2_TLE4_HTR2C
L5_6_FEZF2_TLE4_SCUBE1
L5_6_THEMIS_NR4A2_NTNG2
L5_6_THEMIS_SEMA3A
Neu_dev_THY1
TBD
ID2_CHRNA7_CNR1
ID2_CHRNA7_NCKAP5
ID2_CHRNA7_PAX6
ID2_LAMP5_CRH
ID2_LAMP5_MEIS2
ID2_LAMP5_NMBR
ID2_LAMP5_NOS1
PVALB_BTBD11
PVALB_LGR5
PVALB_MEPE
PVALB_UNC5B
SST_CALB1
SST_ISOC_THSD7B
SST_NOS1
SST_NPY
SST_STK32A
SST_TAC3
SST_TH
VIP_ABI3BP
VIP_CBLN1
VIP_CRH
VIP_NRG1
VIP_SEMA3C
VIP_TRPM3_SSTR1
VIP_TYR
Log2
Fold
Change
3
2
1
0
−1
−2
−3

**Extended Data Fig. 14. Heatmap showing DE of genes from mTOR pathway in TSC neuronal subtypes.** Genes are ranked from the top to the bottom based on how upstream/downstream they are in the mTOR signaling pathway. Genes upstream in the pathway are typically involved in the initial stages of signal transduction, such as those involved in growth factor, nutrient, or energy sensing, while genes downstream are effectors or regulators that are directly influenced by mTOR complex. Orange/blue coloring represents the $log_2$-fold magnitude of gene expression changes.

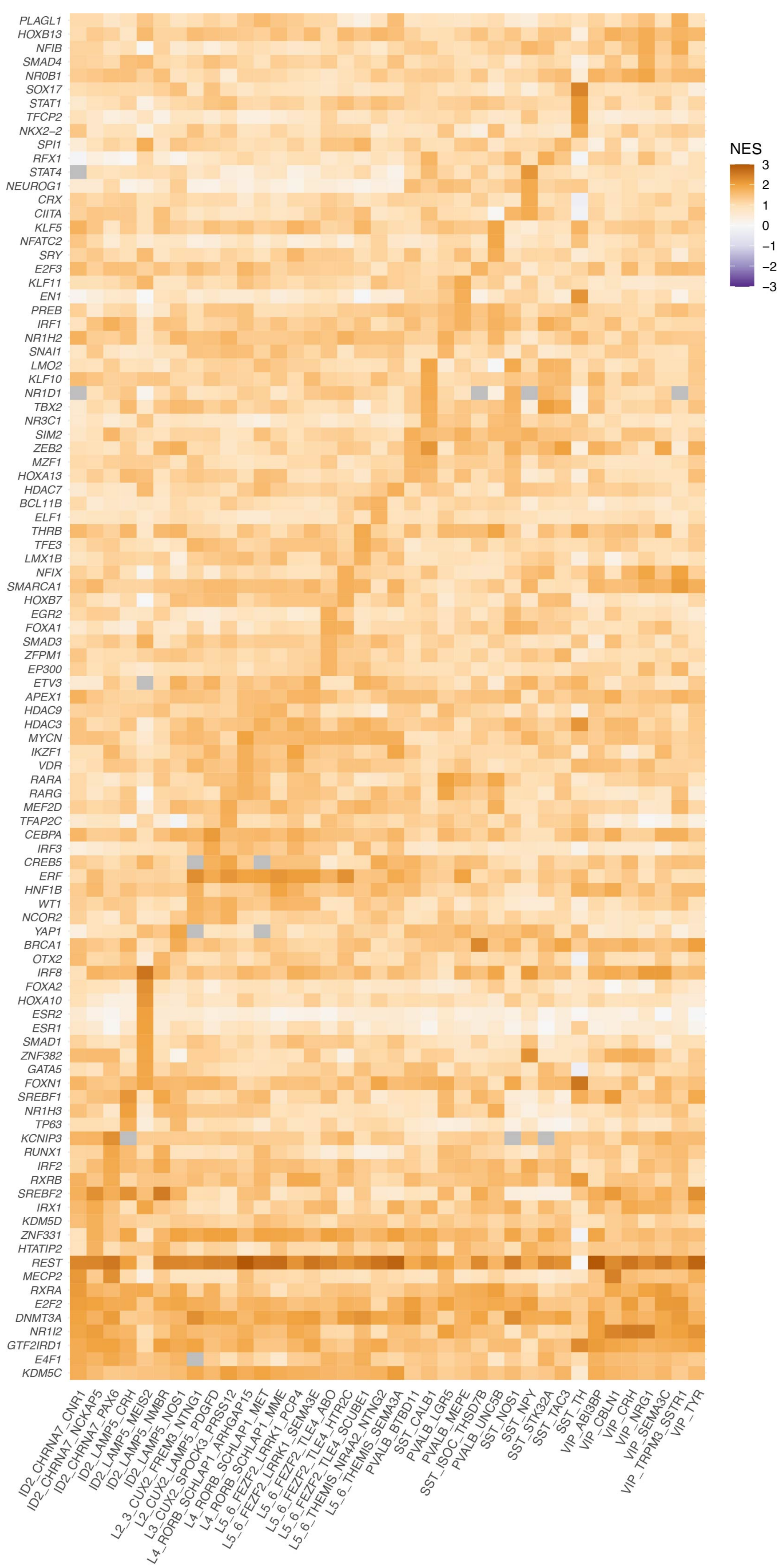
NES
3
2
1
0
−1
−2
−3
PLAGL1
HOXB13
NFIB
SMAD4
NR0B1
SOX17
STAT1
TFCP2
NKX2−2
SPI1
RFX1
STAT4
NEUROG1
CRX
CIITA
KLF5
NFATC2
SRY
E2F3
KLF11
EN1
PREB
IRF1
NR1H2
SNAI1
LMO2
KLF10
NR1D1
TBX2
NR3C1
SIM2
ZEB2
MZF1
HOXA13
HDAC7
BCL11B
ELF1
THRB
TFE3
LMX1B
NFIX
SMARCA1
HOXB7
EGR2
FOXA1
SMAD3
ZFPM1
EP300
ETV3
APEX1
HDAC9
HDAC3
MYCN
IKZF1
VDR
RARA
RARG
MEF2D
TFAP2C
CEBPA
IRF3
CREB5
ERF
HNF1B
WT1
NCOR2
YAP1
BRCA1
OTX2
IRF8
FOXA2
HOXA10
ESR2
ESR1
SMAD1
ZNF382
GATA5
FOXN1
SREBF1
NR1H3
TP63
KCNIP3
RUNX1
IRF2
RXRB
SREBF2
IRX1
KDM5D
ZNF331
HTATIP2
REST
MECP2
RXRA
E2F2
DNMT3A
NR1I2
GTF2IRD1
E4F1
KDM5C
ID2_CHRNA7_CNR1
ID2_CHRNA7_NCKAP5
ID2_CHRNA7_PAX6
ID2_LAMP5_CRH
ID2_LAMP5_MEIS2
ID2_LAMP5_NMBR
ID2_LAMP5_NOS1
L2_3_CUX2_FREM3_NTNG1
L2_CUX2_LAMP5_PDGFD
L3_CUX2_SPOCK3_PRSS12
L4_RORB_SCHLAP1_ARHGAP15
L4_RORB_SCHLAP1_MET
L4_RORB_SCHLAP1_MME
L5_6_FEZF2_LRRK1_PCP4
L5_6_FEZF2_LRRK1_SEMA3E
L5_6_FEZF2_TLE4_ABO
L5_6_FEZF2_TLE4_HTR2C
L5_6_FEZF2_TLE4_SCUBE1
L5_6_THEMIS_NR4A2_NTNG2
L5_6_THEMIS_SEMA3A
PVALB_BTBD11
PVALB_CALB1
PVALB_LGR5
PVALB_MEPE
PVALB_UNC5B
SST_CALB1
SST_ISOC_THSD7B
SST_NOS1
SST_NPY
SST_STK32A
SST_TAC3
SST_TH
VIP_ABI3BP
VIP_CBLN1
VIP_CRH
VIP_NRG1
VIP_SEMA3C
VIP_TRPM3_SSTR1
VIP_TYR

**Extended Data Fig. 15. Heatmap showing enriched transcription factor (TF) networks.** The networks were identified based the z score of DE genes, filtered for top 10 TF per cell type, scaled to -3,3 and presented by hierarchical clustering. Positively enriched TF networks with specificity towards a single type of neurons are shown. For full list, see Extended Data Table 7. NES, Normalized enrichment score.

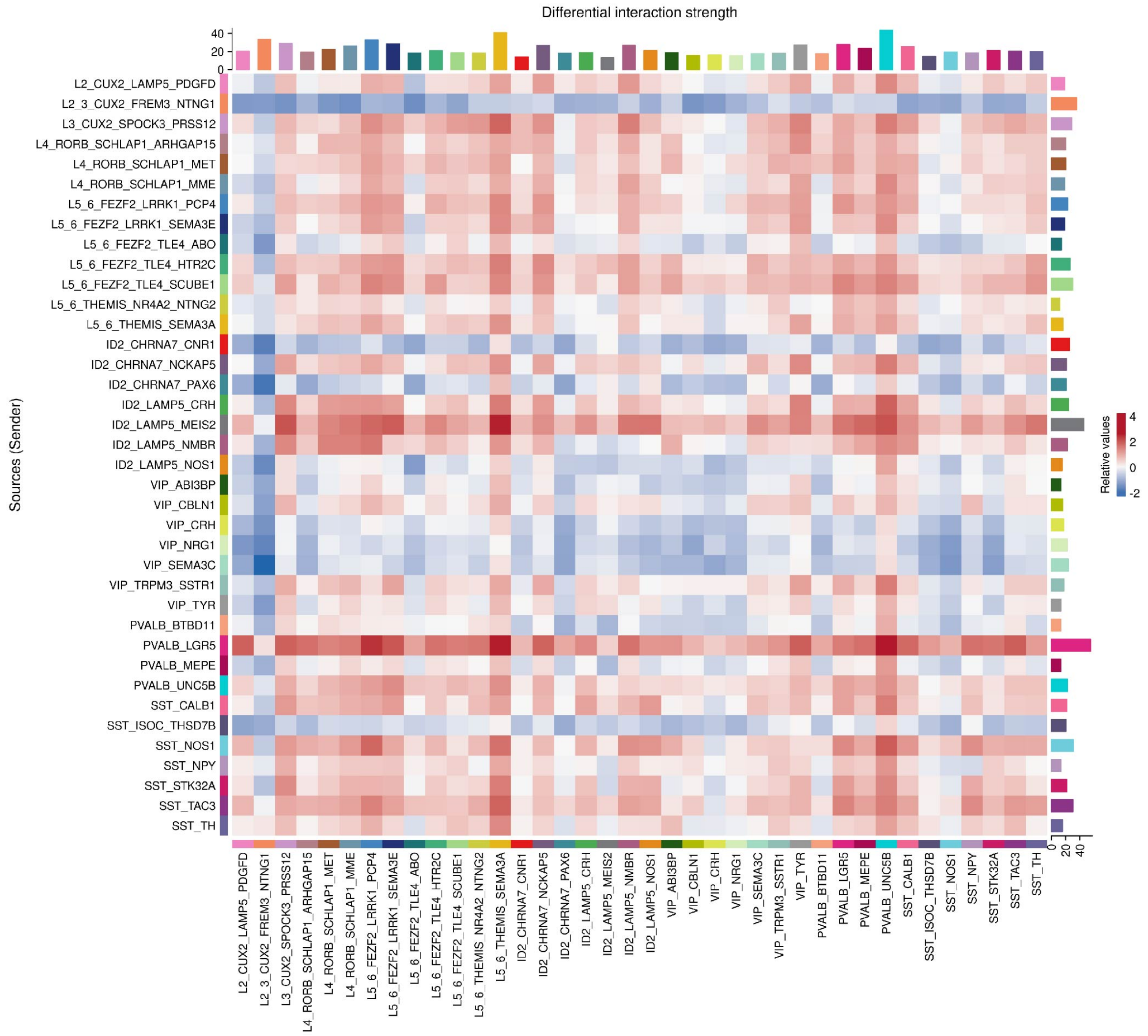


**Extended Data Fig. 16. Heatmap showing differential interaction strength between neuronal cell types.** Sources of signaling are on the y-axis and receivers of the x-axis. The right and top colored bar represents the total weighted interaction strength of outgoing or incoming signaling in absolute values, respectively. Red and blue colors represent increased or decreased signaling strength between neuronal pairs in TSC compared to control, respectively.
